# Neural evidence for boundary updating as the source of the repulsive bias in classification

**DOI:** 10.1101/2023.01.11.523692

**Authors:** Heeseung Lee, Hyang-Jung Lee, Kyoung Whan Choe, Sang-Hun Lee

## Abstract

Binary classification, an act of sorting items into two classes by setting a boundary, is biased by recent history. One common form of such biases is repulsive bias, a tendency to sort an item into the class opposite to its preceding items. Sensory-adaptation and boundary-updating are considered as two contending sources of the repulsive bias, yet no neural support has been provided for either source. Here we explored human brains, using fMRI, to find such supports by relating the brain signals of sensory-adaptation and boundary-updating to human classification behavior. We found that the stimulus-encoding signal in the early visual cortex adapted to previous stimuli, yet its adaptation-related changes were dissociated from current choices. Contrastingly, the boundary-representing signals in the inferior-parietal and superior-temporal cortices shifted to previous stimuli and covaried with current choices. Our exploration points to boundary-updating, rather than sensory-adaptation, as the origin of the repulsive bias in binary classification.

## Introduction

We commit to a proposition about a specific world state when making a perceptual decision. One basic form of such commitments is binary classification. It is to decide whether an item’s magnitude lies on the smaller or larger side of the magnitude distribution across items of interest (**Fig.1A**). For example, when uttering “this tree is tall” while walking in a wood, we are implicitly judging the height of that tree to be taller than the typical height of the trees in the wood^1,2^, where ‘typical height’ works as the boundary dividing the ‘short’ and ‘tall’ classes. Like this, binary classification is exercised in our daily language use—whenever modifying a subject with relative adjectives^3-6^—and has been adopted as an essential paradigm for studying perceptual decision-making^7-17^.

**Fig. 1.**
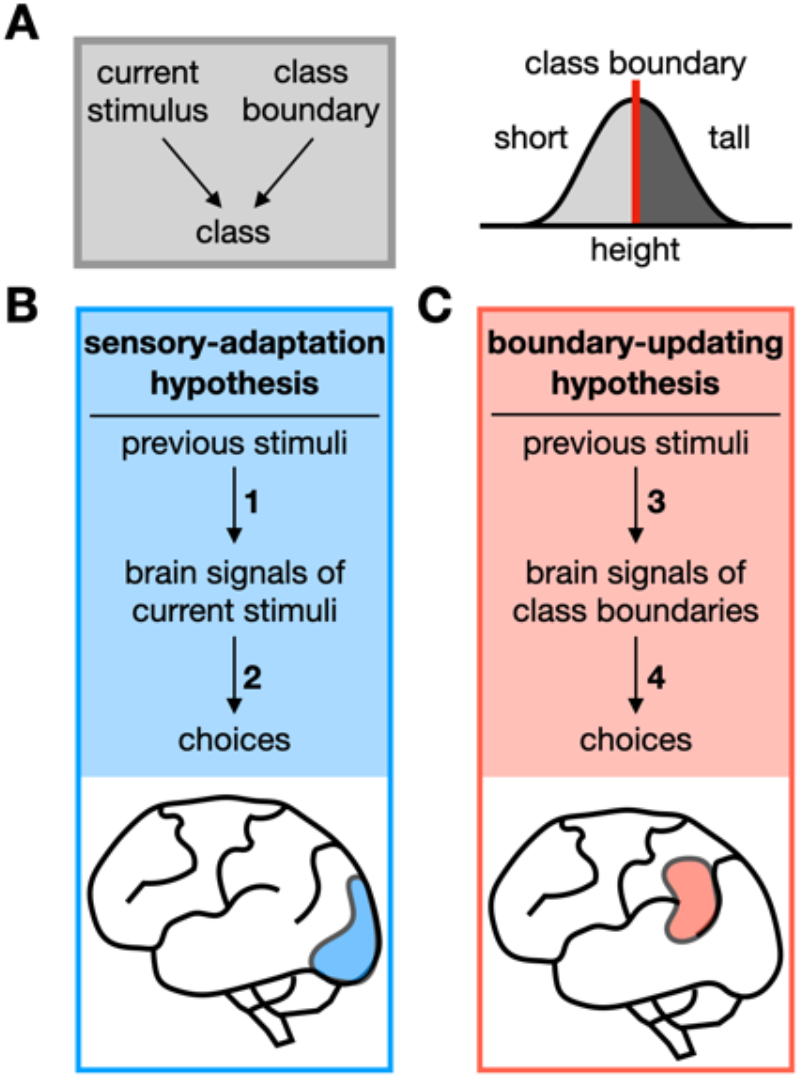
Two contending hypotheses on the origin of the repulsive bias in binary classification. **A** Task structure (left) and statistical knowledge (right) for binary classification. For any given item, its class is determined by its relative position to the class boundary in the distribution of feature magnitudes relevant to a given task (e.g., a tree is classified as ‘tall’ if its height is in the side greater than the typical height of the trees in the wood of interest). This relativity of binary classification makes the ‘biased sensory encoding’ and the ‘biased knowledge about boundary position’ due to previous stimuli, in principle, have equal footings in inducing the repulsive bias. **B** Sensory-adaptation hypothesis. It points to the adaptation of a low-level stimulus-encoding signal to past stimuli (arrow 1) as the origin of the repulsive bias (arrow 2). In the case of visual classification tasks, the task-relevant sensory signals in the early visual cortex (blue patch), which are subject to adaptation, have been hypothesized to mediate the repulsive bias. **C** Boundary-updating hypothesis. It points to the attractive shift of a classifier’s internal class boundary toward previous stimuli (arrow 3) as the origin of the repulsive bias (arrow 4). Such boundary-representing signals are expected to reside not in the early sensory cortex but in the high-tier associative cortices (red patch).

Humans and non-human animals show various forms of history bias in binary classification^17^. One frequent form of such history biases is a tendency to classify an item as the class opposite to its preceding items, dubbed *repulsive bias*^14-17^. For instance, we tend to classify the tree of intermediate height as ‘tall’ after seeing a short tree. Currently, it remains unclear why and how repulsive bias occurs.

As one most straightforward scenario for repulsive bias, the previous stimuli may repel away our perception of the current stimulus from themselves because the sensory system adapts to earlier stimuli^18-24^ (**Fig.1B**). According to this ‘sensory-adaptation’ hypothesis, the current tree is *biasedly* classified as ‘tall’ since the sensory system’s adaptation to the previous short tree makes the current tree appear taller than its physical height. However, there is an alternative scenario, which considers the possibility that the internal class boundary adaptively shifts toward recent samples of property magnitude^14,15,17,25-28^ (**Fig.1C**). According to this ‘boundary-updating’ hypothesis, the current tree is *biasedly* classified as ‘tall’ since the shift of the class boundary toward the previous short tree makes the current tree be positioned in the taller side of the boundary.

As discussed previously^15^, it is hard to assess which hypothesis is more viable based on behavioral data. This hardship arises because binary classification is a matter of the relativity between the perceived stimulus and the class boundary: the identical bias in classification can be caused either by sensory-adaptation or boundary-updating. However, the two hypotheses involve distinct neural routes through which repulsive bias transpires.

The sensory-adaptation hypothesis predicts that the sensory brain signals subject to adaptation—such as those in the early sensory cortex with substantive adaptation to earlier stimuli—contribute to the choice variability. By contrast, the boundary-updating hypothesis predicts that the brain signals of the shifting boundary—such as those in the high-tier cortices involved in the working memory of previous stimuli—contribute to the choice variability.

Here, we tested these two predictions by analyzing functional magnetic resonance imaging (fMRI) data. We found that the stimulus-encoding signal in V1 exhibited adaptation, but its bias induced by adaptation was dissociated from current choices. By contrast, the boundary-representing signals in the posterior-superior-temporal gyrus and the inferior-parietal lobe not only shift to previous stimuli but also covaried with current choices. Our findings contribute to the resolution of the competing ideas regarding the source of repulsive bias by providing the first neural evidence supporting the boundary-updating scenario.

## Results

### Experimental paradigm

Over consecutive trials, participants sorted ring sizes into two classes, *small* and *large*, under moderate time pressure (**Fig.2A**). To ensure decision-makings with uncertainty, we presented three rings (small, medium, and large) differing by a threshold size (Δ), which was tailored for individuals (**Fig.2B; Methods**). The ring sizes were presented in m-sequence to rule out any correlation between consecutive stimulus sizes^30^. We provided participants with feedback after each scan run by summarizing their performance with the proportion of correct trials.

**Fig. 2.**
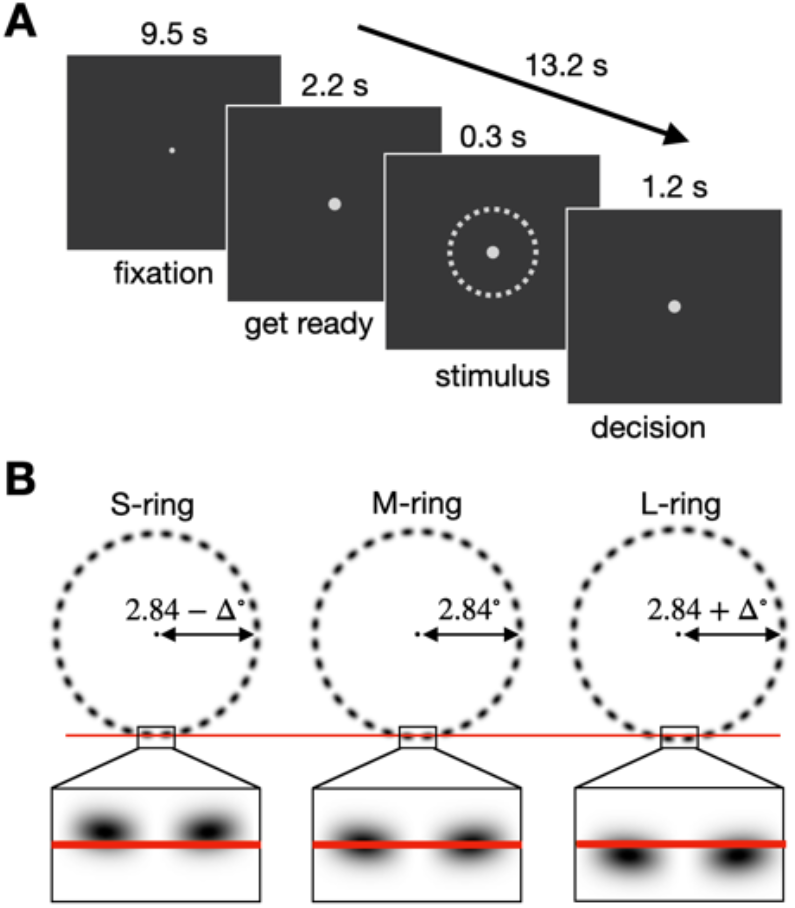
Binary classification task on ring size. **A** Within-trial procedure. With the eyes fixed, human participants were pre-warned, with the increase of the fixation dot, to get ready for the upcoming trial after a long inter-trial interval, briefly viewed the ring stimulus, and judged its size as *large* or *small* in respect to the medium size ring within a limited window of time. **B** Ring stimuli with threshold-level differences in size. On each trial, a participant viewed one of the three rings—small (S), medium (M), large (L), the size contrast (Δ) of which was optimized to ensure threshold-level classification performance on a participant-to-participant basis in a separate calibration run inside the MR scanner, right before the main session of fMRI scan runs. The order of ring sizes over trials was constrained with an m-sequence to preclude the temporal correlation among stimuli. Here, the luminance of the rings is inversed here for an illustrative purpose.

To verify the sensory-adaptation hypothesis, we conducted Experiment 1, where 19 participants performed the classification task while BOLD measurements with a high spatial resolution were acquired only from their early visual cortices. To verify the boundary-updating hypothesis, we conducted Experiment 2, where 18 participants performed the same task while their whole brains were imaged. The data of Experiment 1 had been used in our published work^29^.

### Repulsive bias in Experiment 1

The participants in Experiment 1 displayed a substantive amount of repulsive bias. As anticipated, the proportion of large choices (PL) increased as the ring size on the current trial (*S*_(*t*)_) increased. Importantly, when conditioned on the ring size of the immediately preceding trial (*S*_(*t*−1)_), the psychometric function of PL shifts upward and downward following the S-ring and L-ring trials, respectively (**Fig. 3A**), indicating the presence of repulsive bias.

**Fig. 3.**
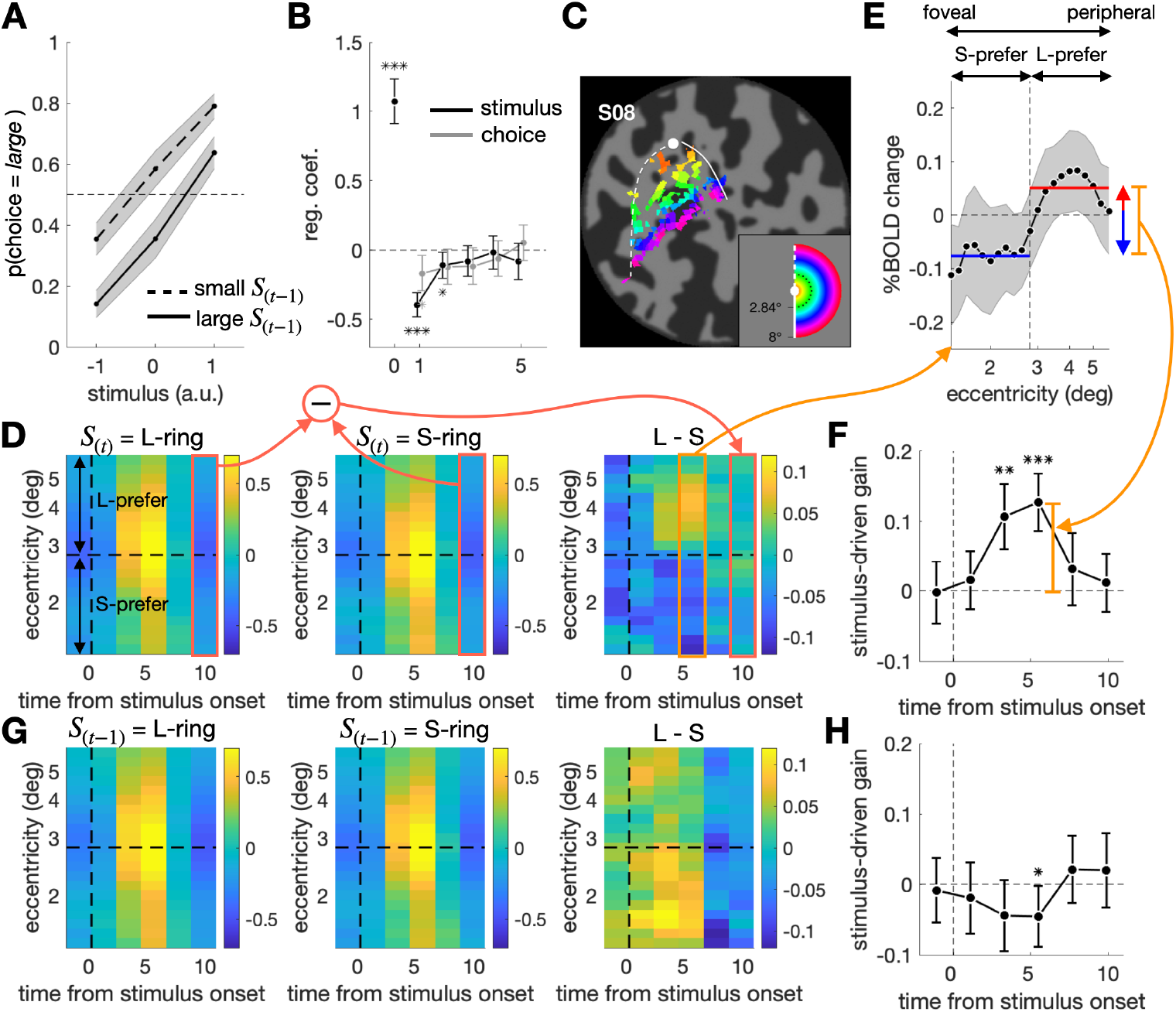
Influences of previous and current stimuli on classification behavior and V1 activity in Experiment 1. **A**,**B** Repulsive bias in psychometric curves (**A**) and regression analysis (**B**). The fraction of *large* choices varied as a function of the current ring size while being modulated concurrently by the previous stimulus (*S*_(*t*−1)_) in a repulsive manner (**A**). The multiple logistic regression coefficients of the current choice were plotted against trial lags (**B**). **C** Eccentricity map of V1 on the flattened left occipital cortex of a representative brain, S08. The dot, curves, and colors correspond to those in the inset depicting the visual field. The image is borrowed from our previous work_29_. **D**,**G** Spatiotemporal BOLD V1 responses to L-ring (left) and S-ring (middle), and their differentials (right), presented on the current (**D**) and previous (**G**) trials. The color bars indicate BOLD changes in the unit of % signal, averaged across all participants. The vertical dashed line marks the time point for stimulus onset. The horizontal dashed line corresponds to the eccentricity of M-ring, splitting the voxels into ‘L-prefer’ and ‘S-prefer’ groups based on their preferred ring size. **E** The differential of BOLD responses at peak between the small and large ring on the current trial. The vertical dashed line marks the eccentricity of M-ring. The horizontal red and blue lines mark the average BOLD signals of the L-prefer and S-prefer voxels, respectively. The vertical orange line quantifies the stimulus-driven gain of V1 responses. **F**,**H** Time courses of the stimulus-driven gain of V1 responses to the current (**F**) and previous (**H**) stimuli. The 95% CIs of the mean across participants are indicated by the shaded areas (**A**,**E**) or by the vertical error bars (**B**,**F**,**H**). Asterisks indicate the statistical significance (*, *P* < 0.05; **, *P* < 10^−8^; ***, *P* < 10^−1^; **B**,**F**,**H**). The orange boxes and arrows are drawn to help the relationships between the panels (**D**,**E**,**F**).

To ensure that this repulsive bias cannot be ascribed to ‘previous decision (*D*_(*t*−1)_)’—owing to its correlation with *S*_(*t*−1)_, we logistically regressed the current choice (*D*_(*t*)_) onto not only *S*_(*t*)_ and *S*_(*t*−1)_ but also *D*_(*t*−1)_. The significant regression coefficients for the previous stimuli (*S*_(*t*−1)_, *β* = −0.39, *P* = 1.6 × 10^−8^; *S*_(*t*−1)_, *β* = −0.11, *P* = 0.026; one sample t-test) confirm the robust presence of repulsive bias in Experiment 1 (**Fig. 3B**).

### Sensory adaptation in V1

As a first step toward the verification of the sensory-adaptation hypothesis, we defined the size-encoding signal in V1. As our group showed previously^29^, the eccentricity-tuned BOLD responses in V1 (**Fig. 3C**) readily resolved the threshold-level differences in ring size, as anticipated by the retinotopic organization of the V1 architecture (**Fig. 3D**). Thus, the subtraction of the BOLD responses at the voxels preferring S-ring to L-ring from those at the voxels preferring L-ring to S-ring (**Fig. 3E**) was significantly greater when *S*_(*t*)_ was large than when small (the third and the fourth time points, *β* = 0.11, *P* = 1.5 × 10^−4^ and *β* = 0.13,*P* = 3.7 × 10^−6^; one sample t-test; **Fig. 3F**).

Next, having defined the size-encoding signal in V1, which will be referred to as ‘*V*1’, we sought the evidence of sensory-adaptation in that signal. According to the previous work on sensory-adaptation^20,31-33^, we expected *V*1 to decrease following the large size and to increase following the small size due to the selective gain reduction at the sensory neurons tuned to previous stimuli. In line with this expectation, *V*1 indeed significantly decreased when preceded by L-ring than when preceded by S-ring (the fourth time point, *β* = −0.45,*P* = 0.040; one sample t-test; **Fig. 3G,H**). In sum, the V1 population activity reliably encoded the ring size and exhibited sensory adaptation.

### The variability of *V*1 associated with previous stimuli fails to contribute to the choice variability

Next, we verified the critical prediction of the sensory-adaptation hypothesis on repulsive bias. In below, we will define what this crucial prediction is and how we empirically examine that prediction.

Above, we confirmed that the ring size, not only on the current trial (*S*_(*t*)_) but also on the previous trial (*S*_(*t*−1)_), affects *V*1 on the current trial (*S*_(*t*−1)_ → *V*1 ← *S*_(*t*)_ in **Fig.4A**). What we do not know yet is whether the variabilities of *V*1 that originate from *S*_(*t*)_ and *S*_(*t*−1)_, respectively, flow all the way into the observer’s current choice (*S*_(*t*)_ → *V*1 → *D*_(*t*)_ and *S*_(*t*−1)_ → *V*1 → *D*_(*t*)_ in **Fig.4A**). Critically, if the sensory-adaptation hypothesis is true, the variability of *V*1 associated with *S*_(*t*−1)_ must contribute to the current choice (*D*_(*t*)_) (*S*_(*t*−1)_ → *V*1 → *D*_(*t*)_). Here, it is important to realize that the mere association between *S*_(*t*)_ and *V*1 (*S*_(*t*)_ → *V*1) does not warrant their contribution to *D*_(*t*)_ (*S*_(*t*)_ → *V*1 → *D*_(*t*)_). Likewise, the association between *S*_(*t*−1)_ and *V*1 (*S*_(*t*−1)_ → *V*1) does not warrant their contribution to *D*_(*t*)_ (*S*_(*t*−1)_ → *V*1 → *D*_(*t*)_).

**Fig. 4.**
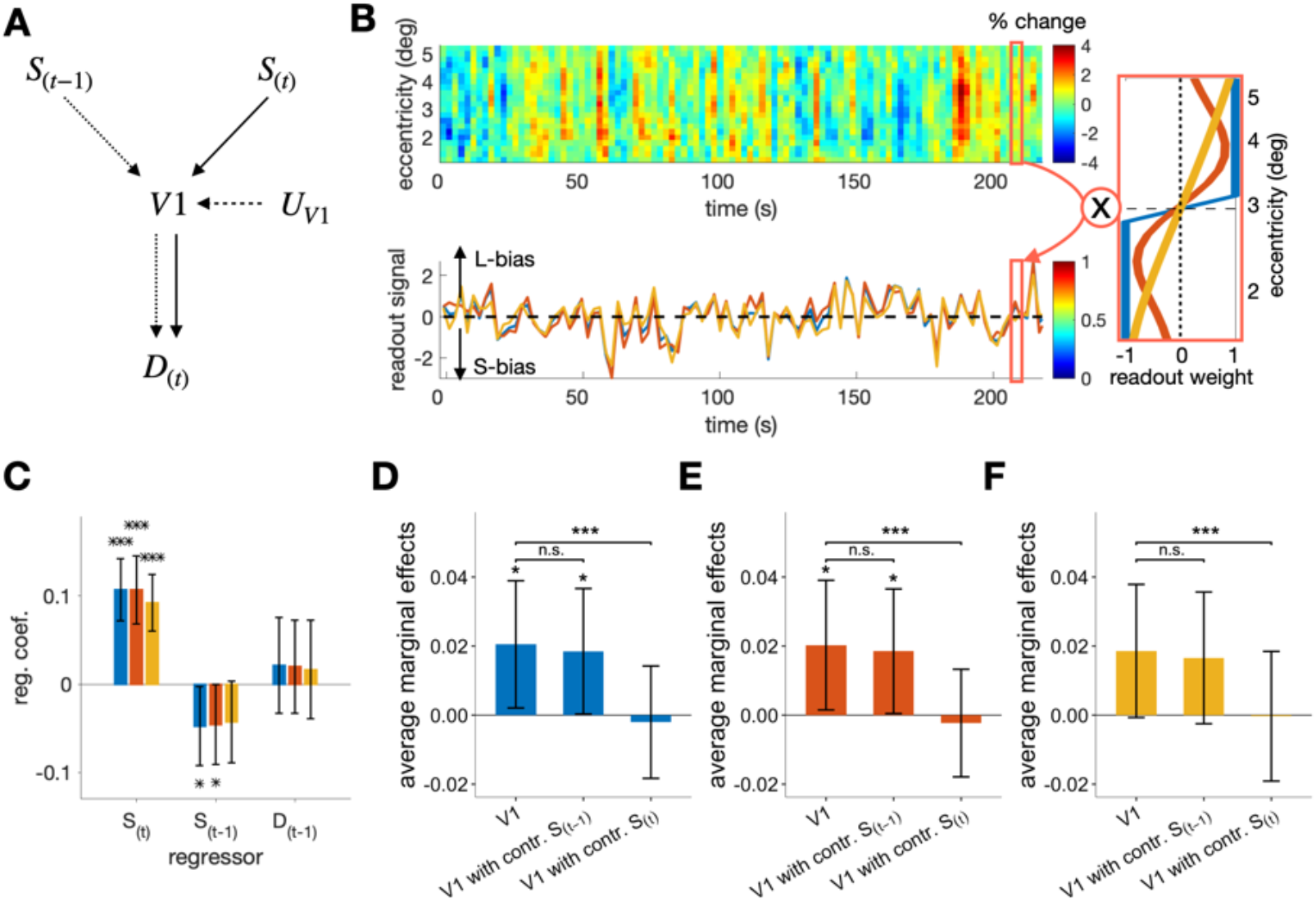
Origin of the covariation between the stimulus-encoding signal of V1 and the current choice. **A** The causal structure of the variables implied by the sensory-adaptation hypothesis. The stimulus-encoding signal of V1 (*V*1) is influenced by the current stimulus (*S*_(*t*)_), the previous stimulus (*S*_(*t*−1)_), and the unknown sources (*U*_*V*−1_). In turn, *V*1 influences the current choice (*D*_(*t*)_). If the sensory-adaptation hypothesis is true, part of the causal influence of *V*1 on *D*_(*t*)_ must originate from *S*_(*t*−1)_, as indicated by the connected chain of the dotted arrows. **B** Extraction of the stimulus-encoding signal of V1. For any given run from any participant, the matrix of spatiotemporal BOLD responses in V1 (top left) was multiplied by one of the three weighting vectors (right; blue, red, and yellow lines represent the uniform, discriminability, and log-likelihood ratio readout schemes, respectively) to result in the vector of stimulus-encoding signal (*V*1) in the same trial length (bottom left). The positive and negative values of *V*1 indicate the larger and smaller sizes of the ring, respectively. **C** Multiple linear regression of the stimulus-encoding signal of V1 on *S*_(*t*)_, *S*_(*t*−1)_, and *D*_(*t*−1)_. The colors correspond to the three different readout schemes in **B. D-F** The average marginal effects (AMEs) of *V*1 on *D*_(*t*)_, with *V*1 extracted by the uniform (**D**), discriminability (**E**), and log-likelihood ratio (**F**) readout schemes. In each panel, the influence of *V*1 on *D*_(*t*)_ that can be ascribed to *S*_(*t*−1)_ and *S*_(*t*)_ were assessed by checking whether the AME of *V*1 on *D*_(*t*)_ (left bar) is significantly reduced or not after controlling the influence of *S*_(*t*−1)_ (center bar) and *S*_(*t*)_ (right bar), respectively. Asterisks indicate the statistical significance (*, *P* < 0.05; **, *P* < 0.01; ***, *P* < 0.001), and “n.s.” stands for the non-significance of the test (**C-F**). The 95% CIs of the mean across participants are indicated by the vertical error bars (**C-F**).

We can test the critical implication of the sensory-adaptation hypothesis by comparing the average marginal effect (AME)^34^ of *V*1 on *D*_(*t*)_ (*V*1 → *D*_(*t*)_) to that of *V*1 on *D*_(*t*)_ with *S*_(*t*−1)_ controlled (*S*_(*t*−1)_ ↛ *V*1 → *D*_(*t*)_). The rationale behind this comparison is that the contribution of *V*1 to *D*_(*t*)_ must be substantially smaller when *S*_(*t*−1)_ was controlled than when not if the contribution of *S*_(*t*−1)_ to *D*_(*t*)_ via *V*1 (i.e., *S*_(*t*−1)_ → *V*1 → *D*_(*t*)_) is substantial. AME was adopted instead of comparing regression coefficients because it does not suffer from the scale problem unlike logistic and probit regression coefficients^35^.

In doing so, the trial-to-trial measures of *V*1 were acquired by taking the sum of BOLDs across the eccentricity bins with the same readout weights used in the previous section (**Fig.4B**). At first, we confirmed that *V*1 contains both current stimuli but also adaptation signals by regressing *V*1 on to *S*_(*t*)_, *S*_(*t*−1)_, and *D*_(*t*−1)_ concurrently (**Fig.4C**). This multiple regression analysis indicates that the previously observed adaptation to *S*_(*t*−1)_ (**Fig.3H**) was still significant (*β* = −0.047, *P* = 0.039), even when we controlled the variability of *D*_(*t*−1)_, a potential confounding variable.

The AME of *V*1 on *D*_(*t*)_ was significant (*β* = 0.020, *P* = 0.031; **Fig.4D**, left). Importantly, it did not decrease when the contribution of *S*_(*t*−1)_ was controlled (*P* = 0.13, one sample t-test; **Fig.4D**, center). Given the significant repulsive bias associated with *S*_(*t*−1)_ presented on the 2-back trial, we also controlled *S*_(*t*−1)_ in addition to *S*_(*t*−1)_. Despite this additional control, the AME of *V*1 on *D*_(*t*)_ did not decrease (*P* = 0.15). These results indicate that the contribution of the previous stimuli to *D*_(*t*)_ via *V*1 is absent or negligible, which is at odds with the sensory-adaptation hypothesis. By contrast, the AME of *V*1 on *D*_(*t*)_ substantively decreased, almost to none, when *S*_(*t*)_ was controlled (*P* = 1.1 × 10^−5^, one sample t-test; **Fig.4D**, right). Put together, the analysis of AME suggests that the contribution of *V*1 to the current choice is ascribed mostly to the current stimulus but hardly to the previous stimuli.

The same pattern of AMEs was observed when we used two alternative readout schemes for extracting *V*1 (The discriminability scheme: *P* = 0.19, *P* = 4.3 × 10^−5^; the log likelihood scheme: *P* = 0.14, *P* = 1.1 × 10^−5^; the one sample t-tests of the AME differences after *S*_(*t*−15_ and *S*_(*t*)_ were controlled, respectively; **Fig.4E,F**).

### Repulsive bias in Experiment 2

Having failed to find the evidence supporting the sensory-adaptation hypothesis in Experiment 1, we conducted Experiment 2 to search the whole brain for the signal representing the class boundary and to test whether that signal relates to the previous stimuli and the current choice in a manner consistent with the boundary-updating hypothesis. As mentioned earlier (see the first Results section), the experimental procedure in Experiment 2 was the same as in Experiment 1, except for the fMRI protocol.

The behavioral performance in Experiment 2 (**Fig.5A,B**) closely matched that in Experiment 1 (**Fig.3A,B**) in many aspects, including the significant presence of repulsive bias (*S*_(*t*−1)_, *β* = −0.54, *P* = 4.6 × 10^−4^; *S*_(*t*−1)_, *β* = −0.24, *P* = 2.3 × 10^−7^; one sample t-test).

**Fig. 5.**
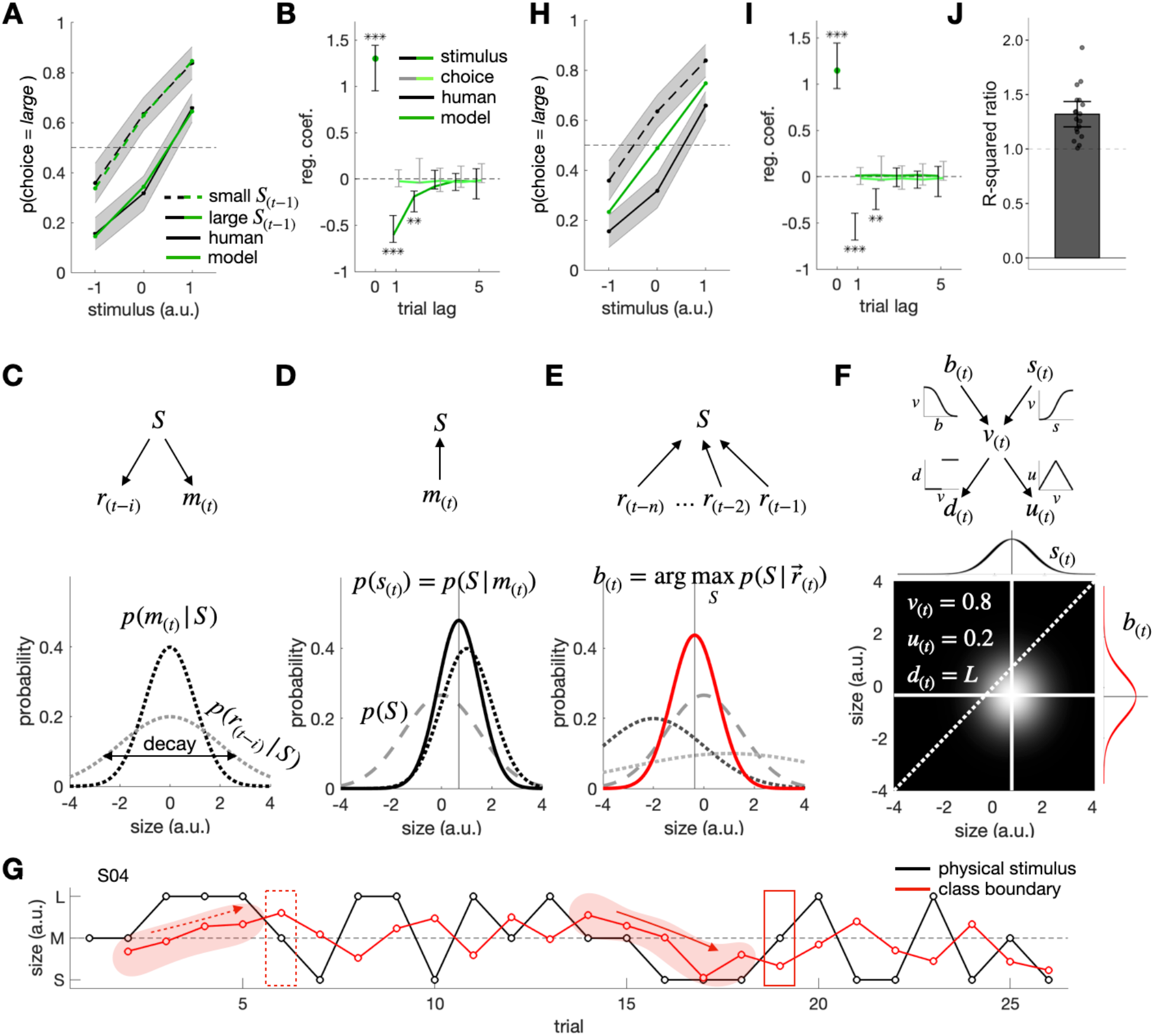
Repulsive bias in Experiment 2 and a Bayesian model of boundary updating (BMBU). **A**,**B** Repulsive bias in psychometric curves (**A**) and regression analysis (**B**). The formats were identical to those in the corresponding figure panels for Experiment 1 (Fig. 3A,B), except that the *ex post* model simulation results (green lines and symbols) are added. **C-F** The measurement generation (**C**), stimulus inference (**D**), class-boundary inference (**E**), and decision-variable deduction (**F**) processes of BMBU. BMBU posits that the Bayesian decision-maker has an internal causal model of how a physical stimulus size (*S*) engenders a current sensory measurement (*m*_(*t*)_) and a retrieved memory measurement from *i*th preceding trial (*r*_(*t*−*i*)_) (**C**, top), which specifies the probability distribution of *m*_(*t*)_ and *r*_(*t*−))_ conditioned on *S*, respectively (**C**, bottom). In turn, *p*(*m*_(*t*)_|*S*) allows the Bayesian decision-maker to infer *S* upon observing *m*_(*t*)_ by combining it with the prior knowledge about *S, p*(*S*), to compute the posterior probability of *S* given *m*_(*t*)_, *p*(*S*|*m*_(*t*)_) (**D**). Similarly, *p*(*r*_(*t*−*i*)_ |*S*) allows for inferring the class boundary (*b*_(*t*)_) upon retrieving the memory of previous sensory measurements 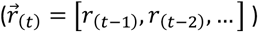 by combining it with *p*(*S*) to compute the posterior probability of *S* given 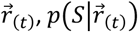 (**E**). In **C-E**, black dotted curves, *p*(*m*_(*t*)_ |*S*); gray dotted curves, *p*(*r*_(*t*−))_1*S*)—the darker the dotted curve is, the more recent the memory is; gray dashed curves, *p*(*S*); black solid curve, *p*(*S*|*m*_(*t*)_); red solid curve, 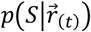. Finally, the inferred stimulus, *s*_(*t*)_, and the inferred class boundary, *b*_(*t*)_, allow for deducing the decision variable, *v*_(*t*)_, the choice variable, *d*_(*t*)_, and the uncertainty variable, *u*_(*t*)_ (**F**, top), as illustrated in an example bivariate distribution of *s*_(*t*)_ and *b*_(*t*)_, from which *v*_(*t*)_, *d*_(*t*)_ and *u*_(*t*)_ are derived (**F**, bottom). **G** An example temporal trajectory of the class boundary inferred by BMBU in a single scan run. The black and red lines indicate the sizes of physical stimulus and the boundary inferred by BMBU, respectively. **H**,**I** *Ex post* simulation results of the constant-boundary model. The formats are identical to those of A and B. **J** The ratio of variance explained by BMBU and the constant-boundary model. Dots represent individual participants.

### Bayesian model of boundary-updating (BMBU)

As we identified *V*1 in Experiment 1, we first need to identify the brain signal that reliably represents the class boundary. However, it is challenging to identify such signals in two aspects. First, unlike in Experiment 1, where V1 was the obvious cortical region to bear the size-encoding signal susceptible to adaptation given a large volume of previous work^22,31-33,36,37^ and our own work^6^, we have no such *a priori* region where the boundary-representing signal resides for sure. This aspect requires us to explore the whole brain. Second, unlike in Experiment 1, where the size variable was physically prescribed by the experimental design, we need to *infer* the trial-to-trial states (i.e., sizes) of the class boundary, which is an unobservable—thus latent—variable. This aspect requires us to build a model. To address these challenges, we inferred the latent state of the class boundary using a Bayesian model of boundary-updating (BMBU) and searched the whole brain for the boundary-representing signal using a searchlight multivariate pattern analysis technique.

We developed BMBU by formalizing the binary classification task in terms of Bayesian decision theory^38^, a powerful framework for modeling human decision-making behavior under uncertainty. Binary classification is to judge whether the ‘ring size on the current trial *t* (*S*_(*t*)_)’ is larger or smaller than the ‘the typical size of the rings appearing across the entire trials 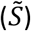.’ Therefore, a classifier must infer them based on the measurements of stimulus size in the sensory and memory systems.

#### The generative model

On trial *t, S*_(*t*)_ is randomly sampled from a probability distribution *p*(*S*) and engenders a measurement in the sensory system *m*_(*t*)_, which is a random sample from a probability distribution *p*(*m*_(*t*)_|*S*_(*t*)_) (black dotted curve of **Fig.5C**). Critically, as *i* trials elapse, *m*_(*t*)_ is re-encoded into a mnemonic measurement in the working-memory system *r*_(*t*−*i*)_, which is a random sample from a probability distribution *p*(*r*_(*t*−*i*)_|*S*_(*t*)_) (light-gray dotted curve in **Fig.5C**). Here, we assumed that the width of *p*(*r*_(*t*−*i*)_|*S*_(*t*)_) increases as *i* increases reflecting the working memory decay^39,40^.

#### Inferring the current stimulus size

On trial *t*, the Bayesian classifier infers *S*_(*t*)_ by inversely propagating *m*_(*t*)_ in the generative model (**Fig.5D**, top). As a result, the inferred size (*s*_(*t*)_) is defined as the value of *S* given *m*_(*t*)_, as captured by the following equation:

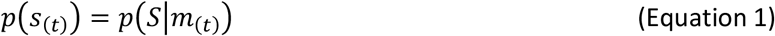

where the width of *p*(*S*|*m*_(*t*)_) reflects the precision of *s*_(*t*)_ (**Fig.5D**, bottom).

#### Inferring the class boundary

On trial *t*, the Bayesian classifier infers the class boundary (*b*_(*t*)_) —i.e., the inferred value of 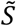—by inversely propagating a set of retrieved measurements in the working memory system 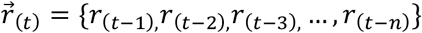 (**Fig.5E**, top). *b*_(*t*)_ is defined as the most probable value of *S* given 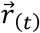, as captured by the following equation:

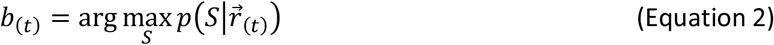

where the width of 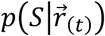 reflects the precision of *b*_(*t*)_. Notably, Equation 2 implies that *b*_(*t*)_ must be attracted more to recent stimuli than to old ones because (i) the imprecision of working memory evidence increases as trials elapse (**Fig.5E**, dotted lines) and (ii) the more uncertain the evidence is, the less weighed the evidence is for class-boundary inference.

#### Making a decision with the inferred current stimulus size and the inferred class boundary

Having estimated *s*_(*t*)_ and *b*_(*t*)_, the Bayesian classifier deduces a decision variable (*v*_(*t*)_) from *s*_(*t*)_ and *b*_(*t*)_ and translating it into a binary decision (*d*_(*t*)_) with a degree of uncertainty (*u*_(*t*)_) (**Fig.5F**). Here, *v*_(*t*)_ is the probability that *s*_(*t*)_ will be greater than *b*_(*t*)_ (*v*_(*t*)_ = *p*(*s*_(*t*)_ > *b*_(*t*)_)); *d*_(*t*)_ is *large* or *small* if *v*_(*t*)_ is greater or smaller than, respectively; *u*_(*t*)_ is the probability that *d*_(*t*)_ will be incorrect (*u*_(*t*)_ = *p*(*s*_(*t*)_ < *b*_(*t*)_|*d*_(*t*)_ = *large*) or *p*(*s*_(*t*)_ > *b*_(*t*)_|*d*_(*t*)_ = *small*))^41^.

In sum, BMBU models a human decision-maker as the Bayesian classifier who, over consecutive trials, continuously infers the class boundary (*b*) and the current stimulus size (*s*), deduces the decision variable (*v*) from *s* and *b*, and makes a decision (*d*) with a varying degree of uncertainty (*u*). As shown below, BMBU well predicts human participants’ choices and reproduces their repulsive bias.

### The prediction and simulation of human choices and repulsive bias by BMBU

We assessed BMBU’s accountability for human behavior in the binary classification task in two aspects, comparing its (i) predictability of the choices and (ii) reproducibility of repulsive bias to those of the control model which does not update the class boundary (‘constant-boundary model’; **Methods**).

We assessed the predictability of BMBU and the constant-boundary model by fitting them to human choices using the maximum likelihood rule (**Methods**). BMBU excels over the constant-boundary model in goodness-of-fit (Δ*AIC* = −10.48, a value well beyond the conventional threshold (−4)^42^: *p* = 0.0195, one sample t-test) and further explained the choice variability (32%, the Nagelkerke R-squared; **Fig. 5J**).

After equipping the models with their best-fit parameters, we assessed their reproducibility by making them simulate the decisions over the same sequence of ring sizes presented to the human participants (**Methods**). From this simulation, we can also vividly appreciate how BMBU updates its class boundary (*b*_(*t*)_) depending on the ring sizes encountered over a sequence of classification trials (**Fig. 5G**). As implied by Equation 2, BMBU continuously shifts *b*_(*t*)_ toward the ring sizes shown in previous trials. Such attractive shifts are pronounced especially when streaks of S-ring (the solid arrow in **Fig. 5G**) or L-ring (the dashed arrow in **Fig. 5G**) appeared over trials. Importantly, we confirmed that such boundary-updating of BMBU reproduces the repulsive bias displayed by the human participants with a remarkable level of resemblance, both for the psychometric curves (**Fig.5A**) and for the coefficients of the stimulus and choice regressors (**Fig.5B**). None of the simulated PLs and coefficients—a total of 17 points—fell outside the 95% confidence intervals of the corresponding human PLs and coefficients. Not surprisingly, the constant-boundary model failed to show any slightest hint of repulsive bias (**Fig.5H,I**). Although we used m-sequences to prevent any auto-correlation among ring sizes, the failure of the constant-boundary model in reproducing repulsive bias reassures that the actual stimulus sequences used in the experiment do not contain any unwanted statistics that might induce spurious kinds of repulsive bias.

In sum, BMBU’s inferences of the class boundary based on past stimuli accounted for a substantive fraction of the choice variability of human classifiers and successfully captured their repulsive bias.

### Brain signals of the class boundary and the other latent variables

With the trial-to-trial states of the latent variables of BMBU, we identified the brain signals of those variables with the following rationale and procedure.

On any given trial *t*, a classifier makes a decision in the manner constrained by the causal structure of BMBU (**Fig.5F**). This causal structure implies two important points to be considered when identifying the neural representations of *b, s* and *v*. First, for any cortical activity, its significant correlation with the variable of interest does not necessarily imply that it represents that variable *per se* but is open to the possibility that it may represent the other variables that are associated with the variable of interest. Second, if any given cortical activity represents the variable of interest, that activity must not violate any of its relationships with the other variables that are implied by the causal structure (**Table 1**; **Methods**).

**Table 1.**
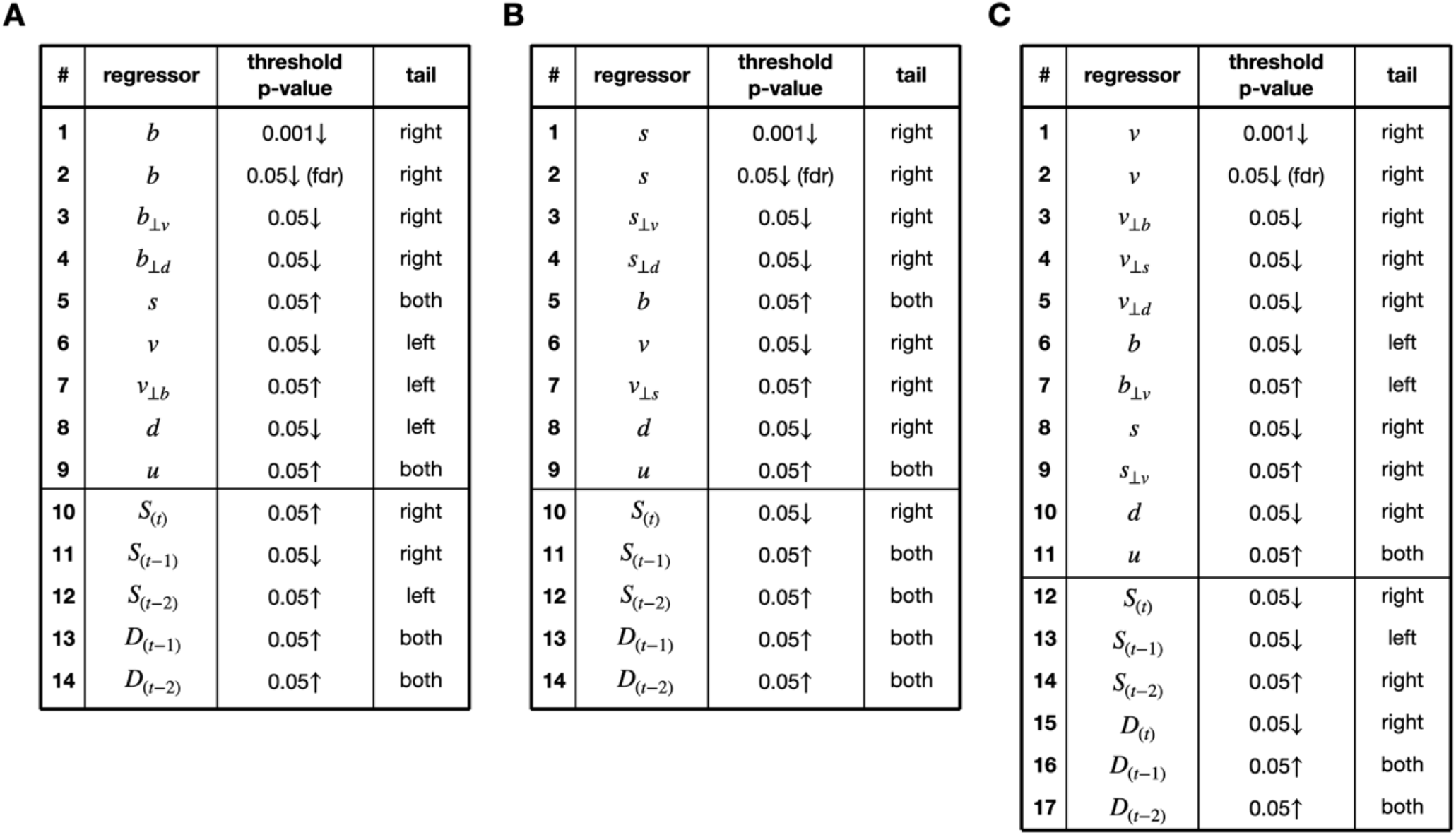
The sets of regressions that BMBU requires the brain signals of its latent variables to satisfy. **A-C** The regressions required for the brain signal of the inferred class boundary (*b*_(*t*)_; **A**), the inferred stimulus (*s*_(*t*)_; **B**), and the decision variable (*v*_(*t*)_; **C**). The top row of each table (#1∼#9 for **A**; #1∼#9 for **B**; #1∼#11 for **C**) specifies the individual, simple regression models in which the brain signal of interest is regressed on a single regressor (second column). Any regressor subscripted with another variable with the perpendicular symbol (e.g., *b*_⊥*v*_) means that the residuals of the left-side variable (e.g., *b*) from the regression of the right-side variable with the perpendicular symbol (e.g., *v*) were used as the regressor. This regression with the *residual regressor* was created to check whether the brain variable of interest has a unique covariation with the original regressor by withholding the influence of the perpendiculared variable (e.g., pSTG_b5_ must be positively correlated with *b* even when the part of *b*’s variability associated with *v* is withheld). The bottom row of each table (#10∼#14 for **A**; #10∼#14 for **B**; #12∼#17 for **C**) specifies the multiple-regression model in which the brain signal of interest is regressed concurrently on the current and previous stimuli and the past or current choices. The third and fourth column of each table specify the statistical criteria used for significance test, where fdr indicates a multiple comparison test controlling the false discovery rate.

| # | regressor | threshold p-value | tail |
| --- | --- | --- | --- |
| 1 | $b$ | 0.001↓ | right |
| 2 | $b$ | 0.05↓ (fdr) | right |
| 3 | $b_{\perp v}$ | 0.05↓ | right |
| 4 | $b_{\perp d}$ | 0.05↓ | right |
| 5 | $s$ | 0.05↑ | both |
| 6 | $v$ | 0.05↓ | left |
| 7 | $v_{\perp b}$ | 0.05↑ | left |
| 8 | $d$ | 0.05↓ | left |
| 9 | $u$ | 0.05↑ | both |
| 10 | $S_{(t)}$ | 0.05↑ | right |
| 11 | $S_{(t-1)}$ | 0.05↓ | right |
| 12 | $S_{(t-2)}$ | 0.05↑ | left |
| 13 | $D_{(t-1)}$ | 0.05↑ | both |
| 14 | $D_{(t-2)}$ | 0.05↑ | both |

B
| # | regressor | threshold p-value | tail |
| --- | --- | --- | --- |
| 1 | $s$ | 0.001↓ | right |
| 2 | $s$ | 0.05↓ (fdr) | right |
| 3 | $s_{\perp v}$ | 0.05↓ | right |
| 4 | $s_{\perp d}$ | 0.05↓ | right |
| 5 | $b$ | 0.05↑ | both |
| 6 | $v$ | 0.05↓ | right |
| 7 | $v_{\perp s}$ | 0.05↑ | right |
| 8 | $d$ | 0.05↓ | right |
| 9 | $u$ | 0.05↑ | both |
| 10 | $S_{(t)}$ | 0.05↓ | right |
| 11 | $S_{(t-1)}$ | 0.05↑ | both |
| 12 | $S_{(t-2)}$ | 0.05↑ | both |
| 13 | $D_{(t-1)}$ | 0.05↑ | both |
| 14 | $D_{(t-2)}$ | 0.05↑ | both |

**Table 1 The sets of regressions that BMBU requires the brain signals of its latent variables to satisfy.**
| # | regressor | threshold p-value | tail |
| --- | --- | --- | --- |
| 1 | $v$ | 0.001↓ | right |
| 2 | $v$ | 0.05↓ (fdr) | right |
| 3 | $v_{\perp b}$ | 0.05↓ | right |
| 4 | $v_{\perp s}$ | 0.05↓ | right |
| 5 | $v_{\perp d}$ | 0.05↓ | right |
| 6 | $b$ | 0.05↓ | left |
| 7 | $b_{\perp v}$ | 0.05↑ | left |
| 8 | $s$ | 0.05↓ | right |
| 9 | $s_{\perp v}$ | 0.05↑ | right |
| 10 | $d$ | 0.05↓ | right |
| 11 | $u$ | 0.05↑ | both |
| 12 | $S_{(t)}$ | 0.05↓ | right |
| 13 | $S_{(t-1)}$ | 0.05↓ | left |
| 14 | $S_{(t-2)}$ | 0.05↑ | right |
| 15 | $D_{(t)}$ | 0.05↓ | right |
| 16 | $D_{(t-1)}$ | 0.05↑ | both |
| 17 | $D_{(t-2)}$ | 0.05↑ | both |

We incorporated these two points in our search of the brain signals of *b, s* and *v*, as follows. Initially, we identified the candidate brain signals of *b, s*, and *v* by localizing the patterns of activities that closely reflect the trial-to-trial states of *b, s*, and *v*. For localization, we used the support vector regressor decoding with the searchlight technique^43,44^, which is highly effective in detecting the local patterns of population fMRI responses associated with the latent variables of computational models^45^. Next, we put those candidate brain signals to a strong test of whether their trial-to-trial states satisfy the causal relationships with the other variables. Specifically, we converted those causal relationships into the empirically testable sets of regression models, respectively for *b* (14 regressions in **Table 1A**), *s* (14 regressions in **Table 1B**) and *v* (17 regressions in **Table 1A**) and checked whether all the regressors’ coefficients derived from the brain signals were consistent with the regression models (**Methods**).

As a result, the brain signals that survived the exhaustive regression tests clustered in six separate regions (**Fig.6A,B**; **Table 2**; **Supplementary Fig.1**). The signal of *b* appeared in three separate regions at different time points relative to stimulus onset, a region in the left inferior parietal lobe at 1.1s (IPL_b1_) and two regions in the left posterior superior temporal gyrus at 3.3 and 5.5s (pSTG_b3_, pSTG_b5_). The signal of *s* appeared in the left dorsolateral prefrontal cortex at 3.3s (DLPFC_s3_) and in the right cerebellum at 5.5s (Cereb_s5_). The signal of *v* appeared in the left anterior superior temporal gyrus at 5.5s (aSTG_v5_).

**Fig. 6.**
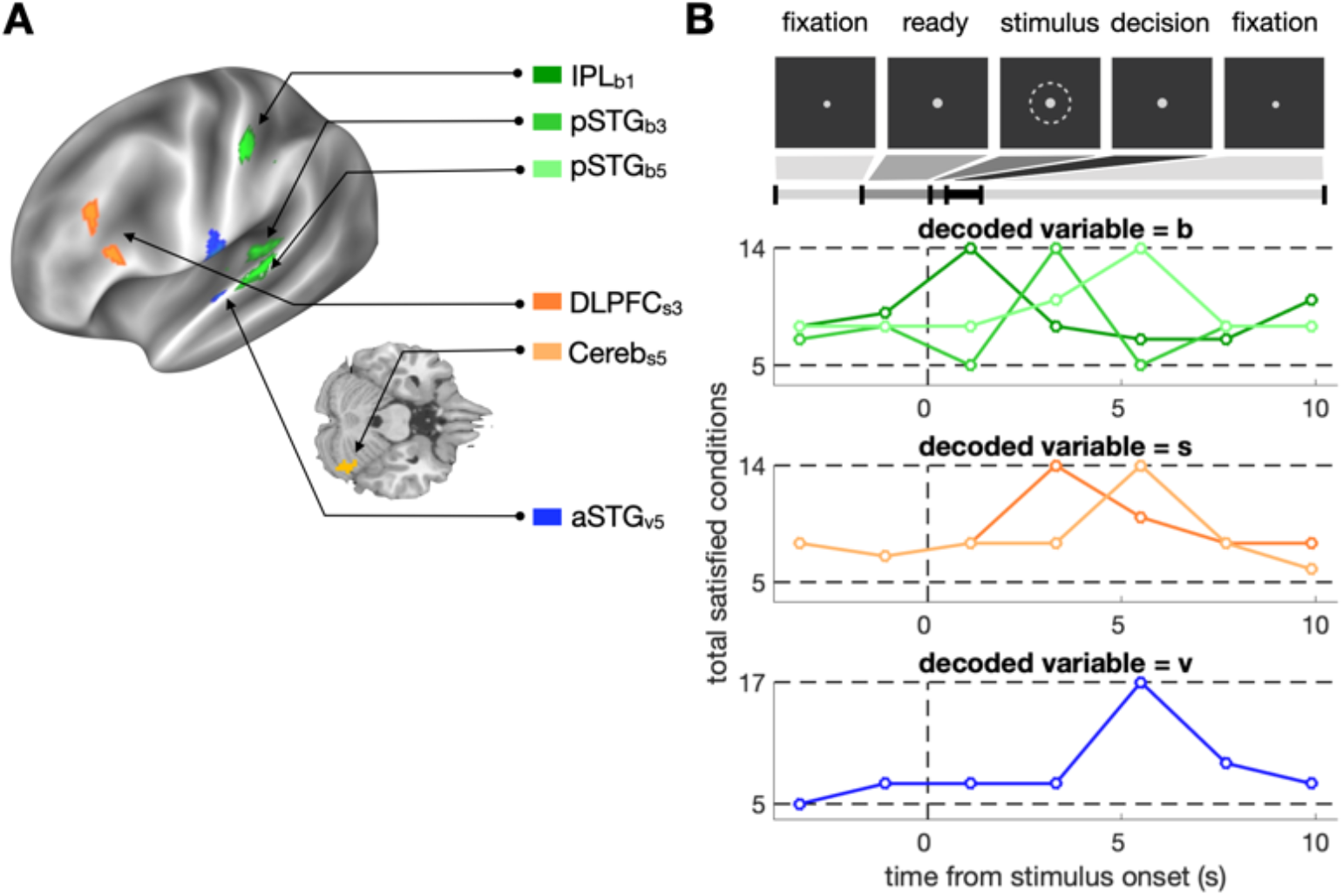
Brain signals of the latent variables of BMBU. **A** Loci of the brain signals. The brain regions where BOLD activity patterns satisfied all the regressions implied by the causal structure of the variables in BMBU are overlaid on the inflated cortex and the axial view of the cerebellum of the template brain. **B** Within-trial time courses of the satisfied regressions in number. The within-trial task phases are displayed (top panel) to help appreciate when the brain signals become pronounced, with the hemodynamic delay (4∼5 s) in BOLD (bottom three panels). The colors of the symbols and lines correspond to those of the brain regions shown in **A**.

**Table 2.** Specification of the brain signals of the latent variables of BMBU.

| abbreviation | cortical area | embodied information | informative time point (s) | contiguous searchlight centers (N) | peak searchlight |  |
| --- | --- | --- | --- | --- | --- | --- |
| | | | | | MNI coordinate | GLMM p-value of $b_{(t)}$ (or $s_{(t)}$, $v_{(t)}$) |
| IPL<sub>b1</sub> | left inferior parietal lobe | $b_{(t)}$ | 1.1 | 15 | [-54, -27, 48] | $8.8 \times 10^{-8}$ |
| pSTG<sub>b3</sub> | left posterior superior temporal gyrus | $b_{(t)}$ | 3.3 | 13 | [-45, -30, 9] | $5.8 \times 10^{-7}$ |
| pSTG<sub>b5</sub> | left posterior superior temporal gyrus | $b_{(t)}$ | 5.5 | 18 | [-66, -21, 9] | $2.3 \times 10^{-7}$ |
| DLPFC<sub>s3</sub> | left dorsolateral prefrontal cortex | $s_{(t)}$ | 3.3 | 33 | [-51, 27, 24] | $1.9 \times 10^{-7}$ |
| Cereb<sub>s5</sub> | right cerebellum | $s_{(t)}$ | 5.5 | 36 | [36, -63, -21] | $1.4 \times 10^{-6}$ |
| aSTG<sub>v5</sub> | left anterior superior temporal gyrus | $v_{(t)}$ | 5.5 | 15 | [-60, -9, 12] | $4.9 \times 10^{-8}$ |

Lastly, for the brain signals of *b, s* and *v* that survived the second step, we confirmed whether the causal structure of *b, s* and *v* prescribed by BMBU is the most likely structure constituted by the brain signals. Specifically, we calculated the Bayesian Information Criterion (BIC) for all the possible causal structures between the brain signals of *b, s* and *v*. The outcomes of BIC evaluation were consistent with BMBU in two aspects. First, out of the 162 possible causal graphs, the smallest (best) BIC value was found for ‘pSTG_b5_→aSTG_v5_ ←Cereb_s5_’ (**Fig. 7**), the one implied by BMBU (**Fig.5F**). Second, note that any graph that includes causal arrows between *b* and *s* would falsify BMBU because BMBU is built upon the assumption that *b* and *s* are independent of one another (i.e., *b* and *s* are biased by previous and current stimuli, respectively) (**Fig.5D,E**). We found that any graph with the causal arrows between *b*_(*t*)_ and *s*_(*t*)_ is significantly less likely than the best causal graph (BIC>2; shown at the bottom of **Fig. 7**)^46^. The results indicate that the relationship between the identified brain signals faithfully reflect the causal relationship of the latent variables implied by BMBU.

**Fig. 7.**
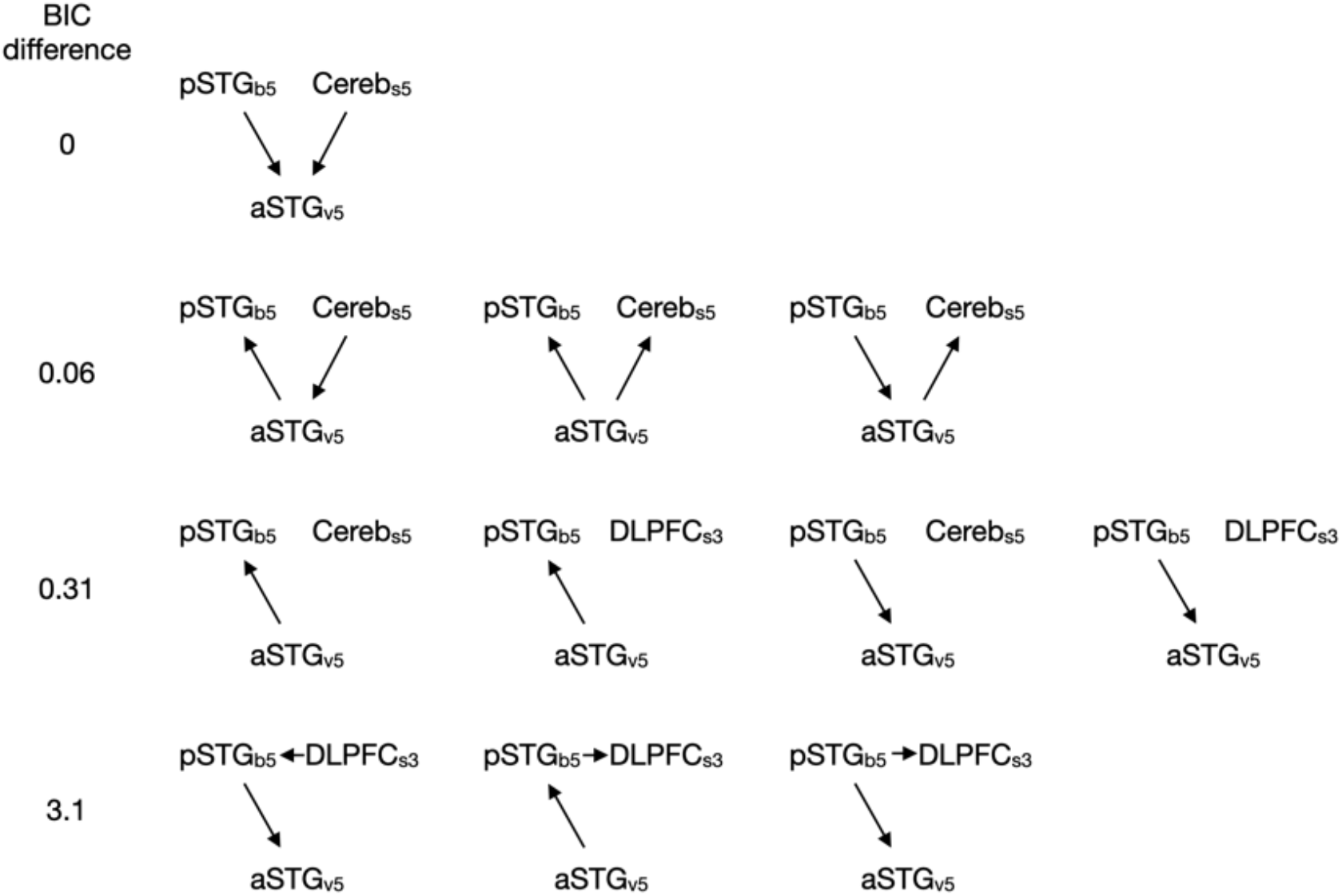
The probable causal structures between the brain signals of the latent variables in BMBU. For each row, the value in the left indicates the relative BIC scores of the causal structures in reference to the most probable one at the top.

### The variability of the class-boundary brain signals associated with previous stimuli contributes to the variability of choice

Finally, with the brain signals that represent the class boundary (IPL_b1,_ pSTG_b3, and_ pSTG_b5_) in our hands, we verified the boundary-updating hypothesis with the rationale and analysis identical to those for the verification of the sensory-adaptation hypothesis.

We stress that the respective associations of the brain signal of *b* with the previous stimulus (*S*_(*t*−1)_; the eleventh row of **Table 1A**) and with the variable *d* (the eighth row of **Table 1A**) do not necessarily imply that the variability of the brain signal of *b* that is associated with *S*_(*t*−1)_ contributes to the choice variability (as implied by the causal information flows through *b* depicted in **Fig.8A**), for the same reasons mentioned when verifying the sensory-adaptation hypothesis. To verify such contribution, we need to compare the AME of the brain signals of *b* on the current choice (*D*_(*t*)_) (pSTG_b5_→ *D*_(*t*)_) to the AME of the brain signals of *b* on *D*_(*t*)_ with *S*_(*t*−1)_ controlled (*S*_(*t*−1)_ ↛pSTG_b5_→ *D*_(*t*)_).

**Fig. 8.**
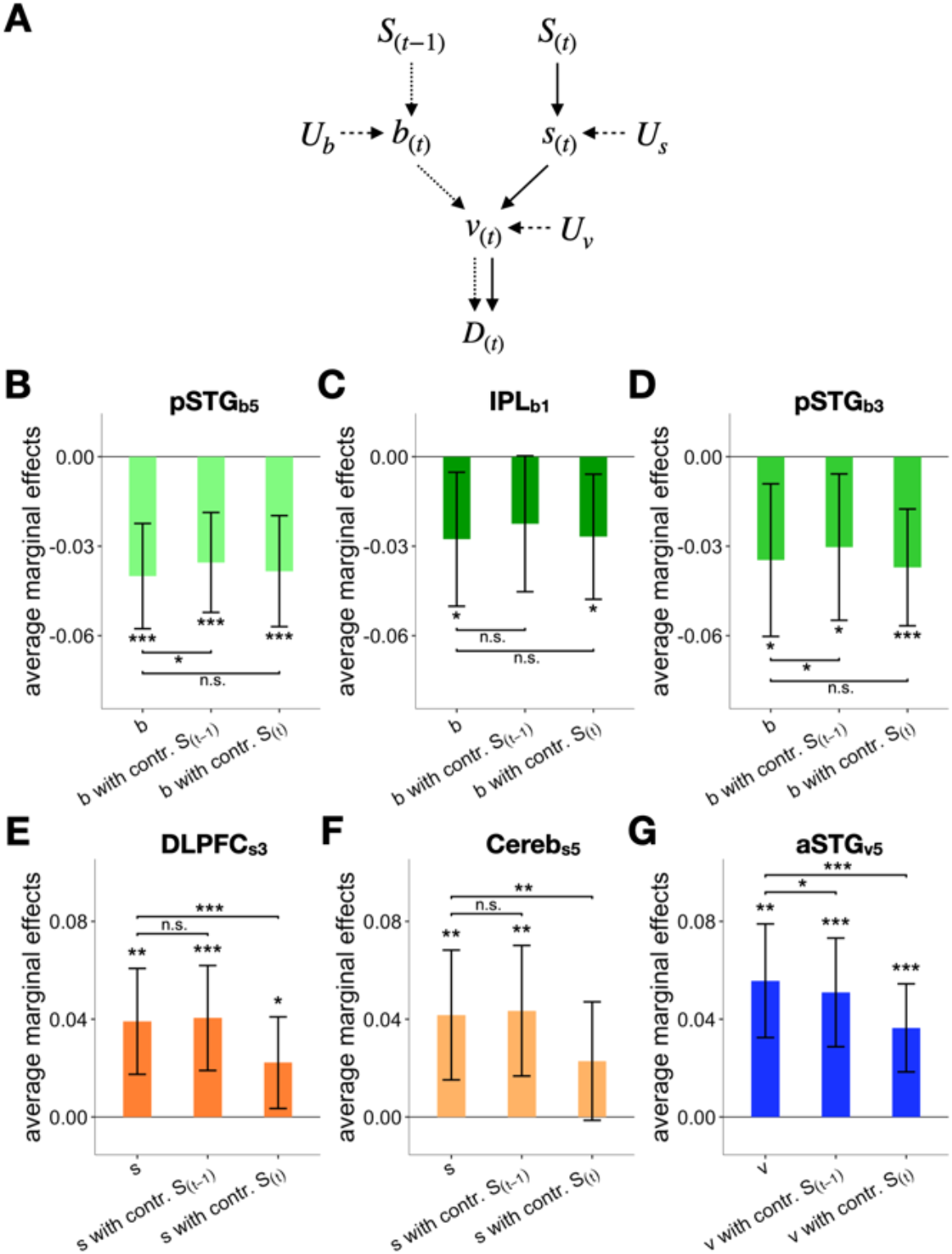
Origin of the covariation between the current choice and the brain signals of the latent variables in BMBU. **A** The causal structure of the variables implied by the boundary-updating hypothesis. The brain signal of the decision variable (*v*_(*t*)_) is influenced by the brain signal of the inferred class criterion (*b*_(*t*)_), brain signal of the inferred stimulus (*s*_(*t*)_), and the unknown sources (*U*_*v*_). In turn, *b*_(*t*)_ is influenced by the previous stimulus (*S*_(*t*#x2212;1_) and the unknown sources (*U*_*v*_) whereas *s*_(*t*)_ is influenced by the current stimulus (*S*_(*t*)_) and the unknown sources (*U*.). Lastly, *v*_(*t*)_ influences the current choice (*D*_(*t*)_). If the boundary-updating hypothesis is true, part of the causal influence of *b*_(*t*)_ on *D*_(*t*)_ must originate from *S*_(*t*−1)_, as indicated by the connected chain of the dotted arrows. **B-G** The average marginal effects (AMEs) of the brain signals on *D*_(*t*)_, with the brain signals of *b*_(*t*)_ from pSTG_b5_ (B), IPL_b1_ (C), and pSTG_b3_ (D), *s*_(*t*)_ from DLPFC_s3_ (E), and Cereb_s5_ (F), and *v*_(*t*)_ from aSTG_v5_ (G). In each panel, the influences of the given brain signal on *D*_(*t*)_ that can be ascribed to *S*_(*t*−1)_ and *S*_(*t*)_ were assessed by checking whether the AME of the given brain signal on *D*_(*t*)_ (left bar) is significantly reduced or not after controlling the influence of *S*_(*t*−1)_ (center bar) and *S*_(*t*)_ (right bar), respectively. The colors of the bars correspond to those of the brain regions shown in Fig. 6A. Asterisks indicate the statistical significance (*, *P* < 0.05; **, *P* < 0.01; ***, *P* < 0.001), and “n.s.” stands for the non-significance of the test. The 95% CIs of the mean across participants are indicated by the vertical error bars.

As anticipated, the AME of pSTG_b5_ on *D*_(*t*)_ was negatively significant (*β* = −0.040, *P* = 1.7 × 10^−(^; **Fig.8B**, left). Importantly, unlike the size-encoding signal in V1, the absolute size of the AME significantly decreased when the contribution of *S*_(*t*−1)_ was controlled (*β* = −0.0046: *P* = 0.012, one sample t-test; **Fig.8B**, center). On the other hand, controlling *S*_(*t*)_ did not affect the AME of pSTG_b5_ on *D*_(*t*)_ at all (*β* = −0.0017: *P* = 0.77, one sample t-test; **Fig.8B**, right) in consistent with the absence of the contribution of *S*_(*t*)_ on *b* (**Fig.5F**). The same patterns were also observed for IPL_b1_ and pSTG_b3_ (**Fig.8C,D**) – the weakened AMEs after controlling *S*_(*t*−1)_ were also observed for pSTG_b3_ (*β* = −0.0043: *P* = 0.046, one sample t-test**; Fig.8D**, center) and marginally for IPL_b1_ (*β* = −0.0052: *P* = 0.0503, one sample t-test; **Fig.8C**, center). Put together, the analysis of AME suggests that the contribution of the class boundary to the current choice is significantly ascribed to the previous stimuli supporting the boundary-updating hypothesis on repulsive bias.

Having found the evidence supporting the boundary-updating hypothesis in the brain signals of *b*, we also carried out the same AME analysis on the brain signals of *s* and *v* below. Given the causal structure of *b, s* and *v*, the validity of the boundary-updating hypothesis will be reinforced if the brain signals of *s* and *v* also turn out acting as fulfilling their causal roles defined by BMBU. According to BMBU, the contribution of *s* to *D*_(*t*)_ must originate not from *S*_(*t*−1)_ but from the *S*_(*t*)_ (the causal route indicated by the solid arrows in **Fig.8A**). In line with this implication, the AME of both DLPFC_s3_ and Cereb_s5_ on *D*_(*t*)_ was significant (*P* < 0.01 for all areas) and significantly decreased after controlling *S*_(*t*)_ (*P* = 4.1 × 10^−4^ for DLPFC_s3_; *P* = 0.0019 for Cereb_s5_, one sample t-test) but not after controlling *S*_(*t*−1)_ (*P* > 0.25 for all areas, one sample t-test) (**Fig.8E,F**). By contrary, the contribution of *v* to *D*_(*t*)_ must originate not only from *S*_(*t*−1)_ but also from *S*_(*t*)_ (**Fig.8A**). In line with this implication, the AME of aSTG_b5_ on *D*_(*t*)_ was significant (*P* = 9.7 × 10^−5^) and significantly decreased both after controlling *S*_(*t*−1)_ (*P* = 0.012, one sample t-test) and after controlling *S*_(*t*)_ (*P* = 7.5 × 10^−(^, one sample t-test) (**Fig.8G**).

In sum, the results suggest that neural signals of *b* and *s* transferred previous and current stimuli to current decisions, respectively, and the neural signal of *v* transferred both previous and current stimuli to current decisions as BMBU implies, which is consistent with the boundary-updating hypothesis.

## Discussion

Here, we explored the two possible origins of repulsive bias, sensory-adaptation vs boundary-updating, in binary classification tasks. Although *V*1 adapted to the previous stimulus, its variability associated with the previous stimulus failed to contribute to the choice variability. By contrast, the variability associated with the previous stimulus in the boundary-representing signals in IPL and pSTG contributed to the choice variability. These results suggest that the repulsive bias in binary classification is likely to arise as the internal class boundary continuously shift toward the previous stimulus.

### Dissociation between sensory-adaptation in V1 and repulsive bias

What makes sensory-adaptation a viable origin of repulsive bias is not its mere presence but its contribution to repulsive bias. The presence of sensory-adaptation in V1 has been firmly established^20,31-33^ and is the necessary premise for the sensory-adaptation hypothesis to work. What matters is whether the trial-to-trial variability of V1 due to such adaptation exerts its influence on the current choice. Such an influence was not observed in our data.

From a general perspective, our findings demonstrate a dissociation between the impact of previous decision-making episodes on the sensory-cortical activity and the contribution of that sensory-cortical activity to decision-making behavior. In this regard, V1 in the current work acts like the binocular-disparity-encoding signal of V2 neurons in a recent single-cell study on monkeys^47^, where, despite the impact of the history on V2 activity, the variability of V2 activity associated with the history failed to contribute to the history effects on decision-making behavior. In a similar vein, our findings also echo the failure of the sensory-adaptation of V1 in influencing the visual orientation estimation in an fMRI study on human participants^48^. There, while sensory-adaptation was evident along the hierarchy of visual areas including V1, V2, V3, V4 and IPS, the history effect of the previous stimulus on the current estimation behavior was opposite to that expected from sensory-adaptation, which suggests that a downstream mechanism compensates for sensory-adaptation. Such a mechanism has also been called for when the single-cell-recording work on monkeys was trying to explain their intriguing adaptation effects found along the visual processing hierarchy^18,49^. For instance, static visual stimuli engendered prolonged—on the order of tens of seconds—adaptation in the lateral geniculate nucleus but the adaptation in V1 paradoxically short-lived —on the order of hundred milliseconds.

### The representations of the class boundary in IPL and pSTG

To account for the repulsive bias in binary classification, previous studies proposed descriptive models based on the common idea that the internal boundary continuously shift towards the previous stimuli^14,15,17,25-28^. However, the neural concomitant of class-boundary updating has been rarely demonstrated.

To our best knowledge, this issue has so far been addressed by one fMRI work^9^, which reported the class-boundary signal in the left inferior temporal pole. However, several aspects of this work make it hard to consider the reported brain signal to represent the class boundary inducing repulsive bias. First, they experimentally manipulated the class boundary in a block-by-block manner. Thus, it is unclear whether the reportedly boundary-representing signal was updated by previous stimuli trial-to-trial, which is required to induce repulsive bias. Second, in their experiments, there was a tight correlation between the class boundary size and the average stimulus size block-by-block. Due to this confounding factor, one cannot rule out the possibility that the reported brain signal reflects the sensory signal associated with the average stimulus size induced by the current stimulus. By contrast, the brain signal of the class boundary in our work is free from these methodological limitations, because it is updated on a trial-to-trial basis and survived the rigorous set of tests including those addressing possible confounding variables (**Table 1**). In this sense, the current work can be considered the first demonstration of the brain signals representing the class boundary that is dynamically updated in such a way that it can account for repulsive bias.

The recent single-cell studies by Hwang et al.^50^ and Akrami et al.^51^ demonstrated that the posterior parietal cortex (PPC) neurons transfer the history information that biases the current choice, which is consistent with the act of the brain signal of class boundary in IPL in our work. Importantly, Hwang et al.’s findings suggest that the class boundary is maintained elsewhere at the time of decision-making since the PPC inactivation before, but not during, the current trial altered the behavioral history effects. Likewise, Akrami et al.’s findings also suggest that the class boundary is stored in other than PPC since intervening PPC diminished the history bias but did not impair the task performance. This call for additional neural locus of sustaining boundary to explain this finding has been incorporated by a circuit model^52^. In sum, these two studies suggest that PPC is not the only neural locus of boundary representation but did not indicate where that representation resides elsewhere. Here, the current work suggests aSTG as the neural locus of boundary representation. One of the functional roles that have been assigned to aSTG, which is to coordinate the spatial reference frame^53-55^, is consistent with the role of the class boundary in binary classification tasks in that it works as a spatial reference against which the current stimulus is compared.

### The representations of *inferred* stimuli in DLPFC and cerebellum

The brain signals of the *inferred* ring size (*s*_(*t*)_) in DLPFC and cerebellum share many features with V1 in that their covariation with the current choice did not decrease after controlling the previous stimulus but decreased after controlling the current stimulus (**Fig.4D-F**; **Fig.8E,F**). This commonality suggests that DLPFC, cerebellum, and V1 alike route the flow of information originating from the current stimulus. Then, what made V1 ineligible for the brain signal of *s*_(*t*)_?

It is notable that BMBU treats *s*_(*t*)_ as the random variable that has the noise variability in addition to being influenced by the physical stimulus (**Fig.8A**). This means that the brain signal of *s*_(*t*)_ is supposed to be associated with the choice even when the current stimulus was controlled because the noise variability can also influence the current choice, as captured by the concept of ‘choice probability’^56^. However, unlike DLPFC and cerebellum, the AME of V1 on the current choice was completely gone after controlling the current stimuli (**Fig.4D-F**) (*P* > 0.75 for all weight schemes), being disqualified as the brain signal of *s*_(*t*)_.

The residence of the inferred—i.e., subjective or perceived—stimulus representation in DLPFC and cerebellum, instead of V1, seems consistent with previous reports. DLPFC and cerebellum have been well known for their critical involvement in visual awareness^57-61^. By contrast, V1 is likely to be involved more in a faithful representation of physical input than its subjective representation^62^, consistent with the previous findings of our group^29,63^.

## Methods

The data of Experiment 1 (Exp1) and Experiment 2 (Exp2) were acquired from 19 (9 females, aged 20–30 years) and 18 (9 females, aged 20–30 years) participants, respectively. Among the participants, 17 of them participated in both experiments. The Research Ethics Committee of Seoul National University approved the experimental procedures. All participants gave informed consent and were naïve to the purpose of the experiments. High-spatial-resolution images were acquired only from the early visual cortex in Exp1 while the images in Exp2 were acquired from the entire brain with a conventional spatial resolution. The 17 people who provided the data for both experiments participated in three to six behavior-only sessions for training and stimulus calibration, one fMRI session for retinotopy, and two experimental fMRI sessions (one for each experiment). The remaining people also completed the behavioral and retinotopy fMRI sessions with the same protocols but participated in only one of the two experiments.

The data from Exp1 had been used for our previous work^29^. The data of Exp2 has never been used in any previous publication. In the current paper, we describe some basic procedures of Exp1. For more details on Exp1, please refer to the original work^29^.

### Experimental setup

MRI data were collected using a 3 Tesla Siemens Tim Trio scanner equipped with a 12-channel Head Matrix coil at the Seoul National University Brain Imaging Center. Stimuli were generated using MATLAB (MathWorks) in conjunction with MGL (http://justingardner.net/mgl) on a Macintosh computer. Observers looked through an angled mirror attached to the head coil to view the stimuli displayed via an LCD projector (Canon XEED SX60) onto a back-projection screen at the end of the magnet bore at a viewing distance of 87 cm, yielding a field of view of 22×17°.

### Behavioral data acquisition

Figure 2 illustrates the experimental procedures. On each trial, the observer initially viewed a small fixation dot (diameter in visual angle, 0.12°; luminance, 321cd/m^2^) appearing at the center of a dark (luminance, 38cd/m^2^) screen. A slight increase in the size of the fixation dot (from 0.12° to 0.18° in diameter), which was readily detected with foveal vision, forewarned the observer of an upcoming presentation of a test stimulus. The test stimulus was a brief (0.3s) presentation of a thin (full-width at half-maximum of a Gaussian envelope, 0.17°), white (321cd/m^2^), dashed (radial frequency, 32cycles/360°) ring that counter-phase-flickered at 10Hz. After each presentation, participants classified the ring size into *small* or *large* using a left-hand or right-hand key, respectively, within 1.5s from stimulus onset. They were instructed to maintain strict fixation on the fixation dot throughout experimental runs. This behavioral task was performed in three different environments: i) the training sessions, ii) the practice runs of trials inside the MR scanner, and iii) the main scan runs inside the MR scanner, in the following order.

In the training sessions, participants practiced the task intensively over several (3 to 6) sessions (about 1,000 trials per session) in a dim room outside the scanner until they reached an asymptotic level of accuracy. Note that we opted to train observers with the stimuli that were much larger than those for the main experiments (mean radius of 9°) to avoid any unwanted perceptual learning effects at low sensory levels and to train participants to learn the task structure of classification.

In the practice runs of trials inside the MR scanner, participants performed 54 practice trials and then 180 threshold-calibration trials while lying in the magnet bore. On each of the threshold-estimation trials in which consecutive trials were apart from one another by 2.7s., one of 20 different-sized rings was presented according to a multiple random staircase procedure (four randomly interleaved 1-up-2-down staircases, two starting from the easiest stimulus and the other two starting from the hardest one) with trial-to-trial feedback based on the class boundary with the radius of 2.84°. A Weibull function was fit to the psychometric curves obtained from the threshold-calibration trials using a maximum-likelihood procedure. From the fitted Weibull function, the threshold difference in size (Δ in **Fig.2B**) associated with a 70.7% correct proportion of responses was estimated. By finding this threshold for each participant, three threshold-level ring sizes were individually tailored as 2.84−Δ° (small-ring), 2.84° (medium-ring), 2.84+Δ° (large-ring).

In the main scan runs, one of these rings with threshold-level differences was presented in the order defined by an m-sequence (base = 3, power = 3; nine S and L-rings and eight M-rings were presented; all scan runs started with two M-rings)^30^ to null the autocorrelation between stimuli. Participants were not informed of the existence of medium-ring. Importantly, participants did not receive trial-to-trial feedback. Instead, only their run-averaged percent correct based on the trials of small-ring and large-ring was shown during a break after each run, to prevent trial-to-trial feedback from evoking any unwanted brain responses associated with rewards^64,65^ or errors^66-68^. Consecutive trials were apart from one another by 13.2s. In the main scan runs of Exp1 and Exp2, observers performed 156 (6 runs X 26 trials) and 208 (8 runs X 26 trials) trials in total, respectively.

### MRI equipment and acquisition

We acquired three types of MRI images. (1) 3D, T1-weighted, whole-brain images were acquired at the beginning of each functional session: MPRAGE; resolution, 1×1×1mm; field of view (FOV), 256mm; repetition time (TR), 1.9s; time for inversion, 700ms; time to echo (TE), 2.36ms; and flip angle (FA), 9°.

(2) 2D, T1-weighted, in-plane images were acquired at the beginning of each functional session. The parameters for the retinotopy-mapping, the V1 mapping, and the whole brain mapping differed slightly as follows (retinotopy, followed by the V1 mapping, and then by the whole brain mapping): MPRAGE; resolution, 1.078×1.078×2.0 mm, 1.083×1.083×2.3 mm 1.08×1.08×3.3 mm; TR, 1.5s; T1, 700ms; TE, 2.79ms; and FA, 9°).
(3) 2D, T2*-weighted, functional images were acquired during each functional session: gradient EPI; TR, 2.7s, 2.2s, 2.2s; TE, 40ms; FA, 77°, 73°, 73°; FOV, 208mm, 207mm, 208mm; image matrix, 104×104, 90×90, 90×90; slice thickness, 1.8mm with 11% gap, 2mm with 15% slice gap, 3mm with 10% space gap; slice, 30, 22, 32 oblique transfers slices; bandwidth, 858Hz/px, 750Hz/px, 790Hz/px; and effective voxel size, 2.0×2.0×1.998mm, 2.3×2.3×2.3mm, 3.25×3.25×3.3mm).

### Retinotopy-mapping protocol

Standard traveling wave methods^69,70^ were used to define V1, to estimate each participant’s hemodynamic impulse response function (HIRF) of V1, and to estimate V1 voxels’ receptive field center and width. High-contrast and flickering (1.33Hz) dartboard patterns were presented either as 0.89°-thick expanding or contracting rings in two scan runs, as 40°-width clockwise or counterclockwise rotating wedges in four runs or in one run as four stationary, 15°-wide wedges forming two bowties centered on the vertical and horizontal meridians. Each scanning run consisted of 9 repetitions of 27s period of stimulation. The fixation behavior during the scans was assured by monitoring participants’ performance on a fixation task, in which they had to detect any reversal in direction of a small dot rotating around the fixation.

### Data preprocessing of V1 images in the retinotopy-mapping session and the main session of Exp1

All functional EPI images were motion-corrected using SPM8 (http://www.fil.ion.ucl.ac.uk/spm)^71,72^ and then co-registered to the high-resolution reference anatomical volume of the same participant’s brain via the high-resolution inplane image^73^. After co-registration, the images of the retinotopy-mapping scan were resliced, but not spatially smoothed, to the spatial dimensions of the main experimental scans. The area V1 was manually defined on the flattened gray matter cortical surface mainly based on the meridian representations, resulting in 825.4±140.7 (mean±SD across observers) voxels. The individual voxels’ time series were divided by their means to convert them from arbitrary intensity units to percentage modulations and were linearly detrended and high-pass filtered^74^ using custom scripts in MATLAB (MathWorks). The cutoff frequency was 0.0185Hz for the retinotopy-mapping session and 0.0076Hz for the main session. The first 10 (of 90; a length of a cycle) and 6 (of 156; a length of a trial) frames of each run of the retinotopy-mapping session and main session, respectively, were discarded to minimize the effect of transient magnetic saturation and allow the hemodynamic response to reach a steady state. The ‘blood-vessel-clamping’ voxels, which show unusually high variances of fMRI responses, were discarded^75,76^; a voxel was classified as ‘blood-vessel-clamping’ if its variance exceeds 10 times of the median variance value of the entire voxels.

### Data preprocessing of whole-brain images in the main session of Exp2

The whole-brain images of the participants in Exp2 were normalized to the MNI template in the following steps: motion correction, co-registration to whole-brain anatomical images via the in-plane images^73^, spike elimination, slice timing correction, resampling to 3×3×3mm voxel size with the SPM DARTEL Toolbox^77^. Spatial smoothing was not applied to avoid the blurring of the patterns of activity. All the procedures were implemented using SPM8 and SPM12 (http://www.fil.ion.ucl.ac.uk.spm)^71,72^, except for spike elimination, for which we used the AFNI toolbox^78^. The first 6 frames of each functional scan, which correspond to the first trial of each run, were discarded to allow the hemodynamic responses to reach a steady state. Then, the normalized BOLD time series at each voxel, each run, and each brain underwent linear detrending, high-pass filtering (0.0076Hz cut-off frequency with a Butterworth filter), conversion into percent-change signals, and correction for non-neural nuisance signals, which was done by regressing out the mean BOLD activity of cerebrospinal fluid (CSF).

The anatomical masks of CSF, white matter, and gray matter were defined by generating the probability tissue maps for individual participants from T1-weighted images, by smoothing those maps to the normalized MNI space using SPM12, and then by averaging them across participants. Finally, the masks were defined as respective groups of voxels whose probabilities exceed 0.5.

Unfortunately, in a few of the sessions, functional images did not cover the entire brain. Especially, the lost part was much larger in one participant’s session than the others including the orbitofrontal cortice and posterior cerebellum. Thus, not to lose too many of voxels for analysis due to this single session, we relaxed the criterion of voxel selection a bit by including the voxels that were shared by more than 16 brains in the normalized MNI space. As a result, some voxels in the temporal pole, ventral orbitofrontal, and posterior cerebellum were excluded from data analysis.

### Estimation of the eccentricities in retinotopic space for V1 voxels

For each V1 voxel in Exp1, its eccentricity (*e*) was defined by fitting a one-dimensional Gaussian function simultaneously to the time-series of fMRI responses to the expanding and contracting ring stimuli in the retinotopy session, which were also used for the definition of V1. The essence of this procedure is as follows (additional details can be found in the original paper^29^).

First, the time series of fMRI were extracted only from a relevant group of voxels with SNR>3 in both of the ring scan runs. Second, an eccentricity-tuning curve (gain over eccentricity, in other words) of a single voxel, *g*(*ε*), was modeled by a Gaussian as a function of the eccentricity in a visuotopic space, *ε*, and it was parameterized by a peak eccentricity, *e*, and a tuning width, *σ*:

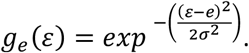

Third, the collective responses of visual neurons within that voxel with a particular *g*(*ε*) at a given time frame *t, n*(*t*), were predicted by multiplying *g*(*ε*) by spatial layout of stimulus input at that time frame, *s*(*ε, t*):

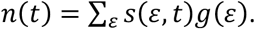

Fourth, the predicted time-series of fMRI responses of that voxel, *fMRI*_*p*_(*t*), were generated by convoluting *n*(*t*) with a scaled (by *β*) copy of the HIRF acquired from the meridian scans, *h*(*t*)*β*, and plus a baseline response, *b*:

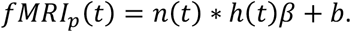

Fifth, the eccentricity *e* and the other model parameters (*σ, β, b*) were found by fitting *fMRI*_*p*_(*t*) to the predicted time-series of fMRI responses to the actual stimulation, *fMRI*_*o*_ (*t*), by minimizing the residual sum of squared errors between *fMRI*_*p*_ (*t*)and *fMRI*_*o*_(*t*) over all time frames, *RSS*:

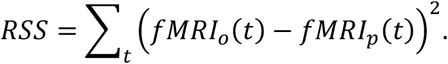

### Extraction of the size-encoding signal from V1 voxels

The three different weighting profiles, each representing the contributions of the individual eccentricity bins assessed by the three different schemes (the uniform, the discriminability, and the log-likelihood ratio schemes), were defined as follows. The uniform scheme (blue in **Fig.4B**) assigned three discrete values to the eccentricity bins depending on which flanking side of the M-ring (*r*_7_) their preferred eccentricities (*e*) belonged to:

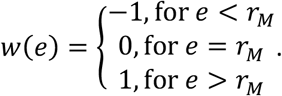

The discriminability scheme (red in **Fig.4B**) defined the weights in proportion to the differential responses of given eccentricity bins to the L (*r*_8_) and the S-rings (*r*_0_), which were derived from the eccentricity-tuning curves defined from the retinotopy-mapping session:

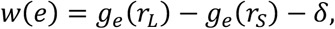

where *g*_1_ is the eccentricity-tuning curve of the eccentricity bin with preferred eccentricity, *e*, and the baseline offset, *δ*, is as follows:

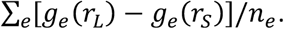

The log-likelihood ratio scheme (yellow in **Fig.4B**) defined the weights by taking the differences between the log-likelihoods of obtaining a given response if the stimulus were the L-ring, *logL*_*L*_, and if the stimulus were the S-ring, *log L*_*S*_. Because the eccentricity-tuning curves were assumed to be described by a Gaussian function, the log-likelihood ratio weights at preferred eccentricity, *e*, can be simplified to the following formula:

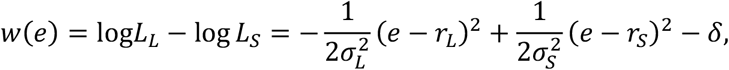

where *σ*_*L*_ and *σ*_*S*_ are the tuning widths with *r*_*L*_ and *r*_*S*_, and the baseline offset, *δ*, is as follows:

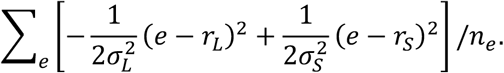

### A Bayesian model of boundary-updating (BMBU)

#### The generative model

The generative model is the observers’ causal account for noisy sensory measurements, where the true ring size, *S*, causes a noisy sensory measurement on a current trial, *m*_(*t*)_, which becomes noisier as *i* trials elapse, thus turning into a noisy retrieved measurement of the value of *S* on trial *t* − *i, r*_(*t*−*i*)_ (**Fig.5C**). Hence, the generative model can be specified with the following three probabilistic terms: a prior of *S, p*(*S*), a likelihood of *S* given *m*_(*t*)_, *p*(*m*_(*t*)_|*S*), and a likelihood of *S* given *r*_(*t*−*i*)_, *p*(*r*_(*t*−*i*)_|*S*). These three terms were all modeled as normal distribution functions, the shape of which is specified with mean and standard deviation parameters, *μ* and *σ*: *μ*_0_ and *σ*_0_ for the prior, 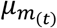 and 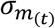 for the likelihood for *m*_(*t*)_, and 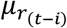 and 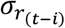 for the likelihood for *r*_(*t*−*i*)_. The mean parameters of the two likelihoods, 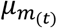 and 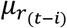, are identical to *m*_(*t*)_ and *r*_(*t*−*i*)_; therefore, the parameters that must be learned are reduced to 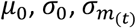, and 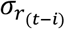.

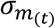 is assumed to be invariant across different values of *m*_(*t*)_, as well as across trials. Therefore, 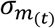 is reduced to a constant *σ*_*m*_. Finally, because 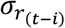 is assumed to originate from *σ*_*m*_ and to increase as trials elapse^39,40^, 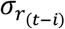 is also reduced to the following parametric function: 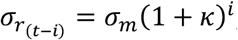, where *κ* > 0. As a result, the generative model is completely specified by the four parameters, Θ = {*μ*_0,_ *σ*_0,_ *σ*_*m*_, *κ*}.

#### Stimulus inference (s)

A Bayesian estimate of the value of *S* on a current trial, *s*_(*t*)_, was distributed as a posterior function of a given sensory measurement *m*_(*t*)_:

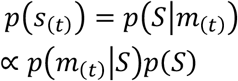

The posterior *p*(*S*|*m*_(*t*)_) is a conjugate normal distribution of the prior and likelihood of *S* given the evidence *m*_(*t*)_ whose mean 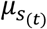 and standard deviation 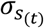 were calculated as follows (**Fig.5D**):

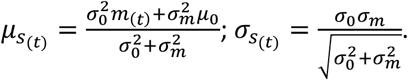

#### Class boundary inference (b)

The Bayesian observer infers the value of class boundary on a current trial, *b*_(*t*)_, by inferring the posterior function of a given set of retrieved sensory measurements 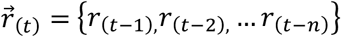:

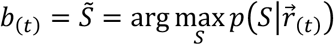

where the maximum number of measurements that can be retrieved, *n*, was set to 7. We set 7 because it is much longer than the effective trial lags of the previous stimulus effect (**Fig.5B**). Here, 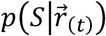 is a conjugate normal distribution of the prior and likelihoods of *S* given the evidence 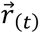:

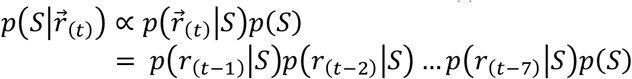

whose mean and standard deviation were calculated^79^ based on the knowledge of how the retrieved stimulus becomes noisier as trials elapse:

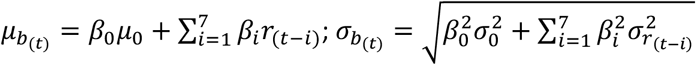

where 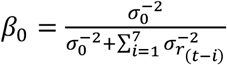 and 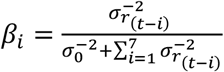. We postulated that the uncertainty of *b* is equivalent to 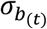 (**Fig.5E,F**).

#### Deduction of decision variable (v), decision (d) and decision uncertainty (u)

On each trial, the Bayesian observer makes a binary decision *d*_(*t*)_ by calculating the probability of *s*_(*t*)_ is larger than *b*_(*t*)_, which is called the decision variable, *v*_(*t*)_, defined as

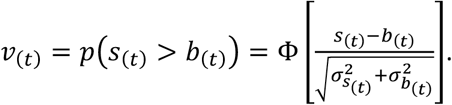

Then, if *v*_(*t*)_ is larger than 0.5, *d*_(*t*)_ is *large*. Otherwise, *d*_(*t*)_ is *small*. Also, we defined the decision uncertainty, *u*_(*t*)_, which represents the odds that the current decision will be incorrect^41^, as follows:

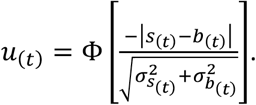

### Fitting the parameters of BMBU

For each human participant, the parameters of the generative model, Θ = {*μ*_9,_ *σ*_9,_ *σ*_*m*_, *κ*}, were estimated as those maximizing the sum of log-likelihoods for *T* individual choices made by the observer, 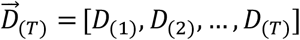

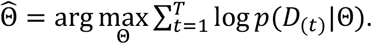

For each participant, estimation was carried out in the following steps. First, we found local minima of parameters using a MATLAB function, *fminseachbnd*.*m*, with the iterative evaluation number set to 50. We repeated this step by choosing 1,000 different initial parameter sets, that were randomly sampled within uniform prior bounds, and acquired 1,000 candidate sets of parameter estimates. Second, from these candidate sets of parameters, we selected the top 20 in terms of goodness-of-fit (sum of log-likelihoods) and searched the minima using each of those 20 sets as initial parameters by increasing the iterative evaluation number to 100,000 and setting tolerances of function and parameters to 10^−7^ for reliable estimation. Finally, using the parameters fitted via the second step, we repeated the second step one more time. Then, we selected the parameter set that showed the largest sum of likelihoods as the final parameter estimates. We discarded i) the first trial of each run and ii) the trials in which RTs were too short (less than 0.3s) for parameter estimation for any further analyses because i) the first trial of each run does not have its previous trial, which is necessary for investigating the repulsive bias, and ii) the response made during the stimulus is shown (0∼0.3s) can be considered too hasty to reflect a normal cognitive decision-making process.

### A constant-boundary model

The constant-boundary model has two parameters, bias of class boundary *μ*_0_ and measurement noise *σ*_*m*_. Stimulus estimates, *s*_(*t*)_, were assumed to be sampled from a normal distribution, 𝒩(*S*_(*t*)_, *σ*_*m*_). Each stimulus sample has uncertainty 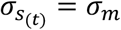. Class boundary *b*_(*t*)_ was assumed to be a constant, *μ*_0_; so 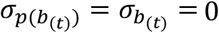.

### Estimation of the latent variables of BMBU

Fitting the model parameters separately for each human participant 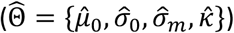 allowed us to create the same number of Bayesian observers, each tailored to each human individual. By repeating the experiment on these Bayesian observers using the stimulus sequences that were identical to those presented to their human partners, we acquired a sufficient number (10^6^ repetitions) of simulated choices, *d*_(*t*)_, and decision uncertainty values, *u*_(*t*)_, which were determined by the corresponding number of the stimulus estimates, *s*_(*t*)_, and the boundary estimates, *b*_(*t*)_, for each Bayesian observer. Then, the averages across those 10^6^ simulations were taken as the final outcomes. When estimating *s*_(*t*)_, *b*_(*t*)_, *v*_(*t*)_, and *u*_(*t*)_ for the observed choice *D*_(*t*)_, we only included the simulation outcomes in which the simulated choice *d*_(*t*)_ matched the observed choice *D*_(*t*)_.

### The multiple logistic regression model for capturing the repulsive bias

To capture the repulsive bias in human classification, we logistically regressed the current choice onto stimuli and choices using the following regression model to obtain regression coefficients 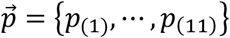 for each observer:

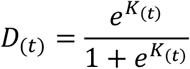

where 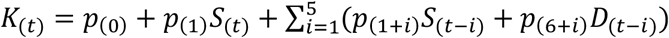, the independent variables were each standardized to z-scores for each participant. *S*_(*t*)_ and *D*_(*t*)_ are the stimulus and the observed choice values at trial *t. S*_(*t*−*i*)_ and *D*_(*t*−*i*)_ are the stimulus and the observed choice at the *i*th trial lags from trial *t*.

To capture the repulsive bias of the Bayesian observers, the Bayesian observers’ choices were also regressed with the logistic regression model by substituting *d*_(*t*)_ and *d*_(*t*−*i*)_, the simulated choices, for *D*_(*t*)_ and *D*_(*t*−*i*)_, the observed choices. The regression was repeatedly carried out for each simulation, and the regression coefficients that were averaged across simulations were taken as final outcomes. The simulation was repeated 10^5^ times. We confirmed that the simulation number was sufficiently large to produce stable simulation outcomes.

### The average marginal effect analysis

Average marginal effect (AME) was calculated by using the R-package ‘margins’^80^. AME quantifies the average marginal effect between an ordinal dependent variable (i.e., binary choice) and an independent variable of a multiple logistic (or probit) regression model^34^. To calculate the AMEs of any given variable on the current choice (*D*_(*t*)_) without controlling the previous (*S*_(*t*−1)_) and current stimuli (*S*_(*t*)_) (i.e., the baseline AME), we implemented a logistic regression model with two regressors –the variable of interest *X* (i.e., V1, *b, s*, or *v*) and the previous choice (*D*_(*t*−1)_):

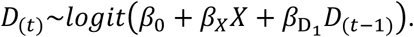

We always included *D*_(*t*−1)_ as a regressor because the effect of *D*_(*t*−1)_ would confound the effect of *S*_(*t*−1)_, if *D*_(*t*−1)_ is not included in the regression model. Specifically, because *S*_(*t*−1)_ and *D*_(*t*−1)_ are highly correlated, it would be unclear whether the AME difference before and after controlling *S*_(*t*−1)_ is ascribed to the effect of *S*_(*t*−1)_ or that of *D*_(*t*−1)_, if *D*_(*t*−1)_ is not controlled. The effect of *D*_(*t*−1)_ was controlled in all regression models.

To test whether the AME of *X* decreased after controlling *S*_(*t*−1)_ (or S_(*t*)_), we calculated the AME of *X* from the logistic regression model including *S*_(*t*−1)_ (or *S*_(*t*)_) as an additional regressor, as follows:

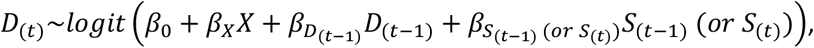

and subtracted the new AME from the baseline AME to see whether the baseline AME significantly changed after controlling previous or current stimuli.

### Searching for the multivoxel patterns of activity representing the latent variables of BMBU

To identify the brain signals of the latent variables of BMBU in fMRI responses, the time-resolved support vector regression (SVR) was carried out in conjunction with a searchlight technique^81,82^. A searchlight has a radius of 9mm (= 3 voxels)^83^ and thus can contain 123 voxels at most. Of the 123 voxels, we excluded the voxels located in CSF or white matter because they reflect non-neural signals. Thus, the effective number of voxels in a searchlight used for the analysis can vary searchlight by searchlight.

We implemented the time-resolved decoding technique in which a target variable is decoded from the BOLD responses at each of the within-trial time points (**Fig.6B**). We used the first four time points (out of six in total) because the BOLD responses associated with the action of button press—the last process of the sensory-to-motor decision-making stream— is maximized at the fourth time point (the result is not shown here). Before training SVR in a searchlight, the BOLD responses in a searchlight and a target variable were z-scored across trials. Then, the z-scored variable was decoded for each searchlight using the cross-validation method of leave-one-run-out (8-fold cross-validation). As a result, for each searchlight and at each time point, we acquired a set of decoded latent variables in all trials. In other words, on each time point, we acquired the 4-dimensional map of the decoded variable (i.e., 3 spatial dimensions and 1 trial dimension).

To calculate the significance of the decoded variable, the 3D spatial maps of the decoded variables were smoothed with a 5mm FWHM Gaussian kernel on each trial. We then regressed the smoothed decoded variable onto the regression conditions of the target variable. The number of regression conditions was 14, 14, and 17 for *b*_(*t*)_, *s*_(*t*)_, and *v*_(*t*)_, respectively (**Table 1**). Those regression models were deduced from the causal structure between the variables of BMBU (see the next section). We accepted a given cluster as the brain signals of *b*_(*t*)_, *s*_(*t*)_, or *v*_(*t*)_ only when they satisfied those regression models over more than 12 contiguous searchlights. For the ROI analysis, the decoded values of a given latent variable were averaged over all searchlights within each ROI.

SVR was conducted using LIBSVM (http://www.csie.ntu.edu.tx/~sjlin/libsvm) with a linear kernel and constant regularization parameter of 1^81,83^. The brain imaging results were visualized using Connectome Workbench^84^ and xjview.

### The regression-model test for verifying the brain signals of *b*_(*t*)_, *s*_(*t*)_, and *v*_(*t*)_

To identify the brain signals of *b*_(*t*)_, *s*_(*t*)_, and *v*_(*t*)_, we defined three respective lists of regressions that must be satisfied by the brain signals. We stress that each of these lists consists of the necessary conditions to be satisfied because the conditions are deduced from the causal structure of the variables in BMBU (**Fig.5F**). Below, we specify the specific regressions that constitute these lists.

The 14 regressions for the brain signal of *b* (**Table 1A**): (*b*1-4), *y*_*b*_, *b* decoded from brain signals, must be regressed onto *b*—the variable it represents—even when the false discovery rate is controlled^85^, and *b* is orthogonalized to *v* or *d*, because it should reflect the variance irreducible to the offspring variables of *b*; (*b*5), *y*_*b*_ must not be regressed onto *s* because *b* and *s* are independent (*b* ↮ *s* **Fig.5F**); (*b*6,7), *y*_*b*_ must be regressed onto *v* (*b* → *v* **Fig.5F**) but not when *v* is orthogonalized to *b* because the influence of *b* on *v* is removed; (*b*8,9) *y*_*b*_ must be regressed onto *d* (*b* → *v* → *d* **Fig.5F**) but not onto *u* because *u* cannot be linearly correlated with *b* (*b* → *v* → *u* is blocked by the interaction between *u* and *v* **Fig.5F**); (*b*10-12), *y*_*b*_ must be regressed onto, not the current stimulus, but the past stimuli—strongly onto the 1-back stimulus and more weakly onto the 2-back stimulus (thus, non-significant regression with one-tailed regression in the opposite sign is modeled conservatively); (*b*13,14), *y*_*b*_ must not be regressed onto previous decisions at all because what updates *b* is not previous decisions but previous stimuli. *b*10-14 were investigated by a multiple regression with regressors [*S*_(*t*)_, *S*_(*t*−1)_, *S*_(*t*−1)_, *D*_(*t*−1)_, *D*_(*t*−1)_]. We did not include *D*_(*t*)_ as a regressor because *D*_(*t*)_ may induce a spurious correlation between *b* and *s* by controlling the collider *v*^86^ (*b* → *v* ← *s* and *v* → *d* **Fig.5F**).

The 14 regressions for the brain signal of *s* (**Table 1B**): (*s*1-4), *y*_=_, *s* decoded from brain signals, must be regressed onto *s*—the variable it represents—even when the false discovery rate is controlled^85^, and *s* is orthogonalized to *v* or *d* because it should reflect the variance irreducible to the offspring variables of *s*; (*s*5), *y*_=_ must not be regressed onto *b* because *s* and *b* are independent of each other (*b* ↮ *s* **Fig.5F**); (*s*6,7), *y*_=_ must be regressed onto *v* (*s* → *v* **Fig.5F**) but not when *v* is orthogonalized to *s* because the influence of *s* on *v* is removed; (*s*8,9) *y*_=_ must be regressed onto *d* (*s* → *v* → *d* **Fig.5F**) but not onto *u* because *u* cannot be linearly correlated with *s* (*s* → *v* → *u* is blocked by the interaction between *u* and *v* **Fig.5F**); (*s*10-12), *y*_=_ must be regressed onto the current stimuli and not the past stimuli because *s* is inferred solely from the current stimulus measurement; (*s*13,14), *y*_=_ must not be regressed onto previous decisions because *s* is inferred solely from the current stimulus measurement. *s*10-14 were investigated by a multiple regression with regressors [*S*_(*t*)_, *S*_(*t*−1)_, *S*_(*t*−1)_, *D*_(*t*−1)_, *D*_(*t*−1)_]. We did not include *D*_(*t*)_ as a regressor because *D*_(*t*)_ may induce a spurious correlation between *b* and *s* by controlling the collider *v*^86^ (*b* → *v* ← *s* and *v* → *d* **Fig.5F**).

The 17 regressions for the brain signal of *v* (**Table 1C**). (*v*1-5), *y*_*v*_, *v* decoded from brain signals, must be regressed onto *v*—the variable it represents—even when the false discovery rate is controlled^85^, and *v* is orthogonalized to *b, s*, or *d*, because it should reflect the variance irreducible to the offspring variables of *v*; (*v*6,7), *y*_*v*_ must be regressed onto one of its parents *b* (*b* → *v* **Fig.5F**), but not when *b* is orthogonalized to *v*, because the influence of *b* on *v* is removed; (*v*8,9), *y*_*v*_ must be regressed onto one of another parent *s* (*s* → *v* **Fig.5F**), but not when *s* is orthogonalized to *v*, because the influence of *s* on *v* is removed; (*v*10,11), *y*_*v*_ must be regressed onto *d* but not onto *u* because *u*’s correlation with its parent *v* cannot be revealed without holding the variability of *d* (the interaction between *u* and *v*); (*v*12-14), *y*_*v*_ must be positively regressed onto the current stimulus because the influence of the current stimulus on *v* is propagated via *s* (*S*_(*t*)_ → *s* → *v*), and negatively regressed onto the past stimuli because the influence of the past stimuli on *v* is propagated via *b* (*S*_(*t*−1)_ → *b* → *v*) —strongly onto the 1-back stimulus and more weakly onto the 2-back stimulus (thus, non-significant regression with one-tailed regression in the opposite sign is modeled moderately); (*v*15-17), *y*_*v*_ must be regressed onto the current decision and not the past decisions because the current decision is a dichotomous translation of *v* (*v* → *d* **Fig.5F**), whereas past decisions have nothing to do with the current state of *v. v*12-17 were investigated by a multiple regression with regressors [*S*_(*t*)_, *S*_(*t*−1)_, *S*_(*t*−1)_, *D*_(*t*)_, *D*_(*t*−1)_, *D*_(*t*−1)_]. *D*_(*t*)_ was included as a regressor because *v* does not suffer from a spurious correlation that arises by controlling a collider variable which is absent in this case.

### Bayesian network analysis

For the data-driven Bayesian network analysis, we derived an exhaustive set of causal graphs between the brain signals of the latent variables of BMBU and calculated BIC for each graph^87^. Specifically, to see whether the causal relationship between decoded *b, s* and *v* is consistent with the causal structure that is postulated by BMBU, we searched for the causal graph (*G*) whose likelihood is maximal given the time series of three brain signals {*y*_*c*_, *y*_*s*_, *y*_*v*_}.

A total of 27 edge structures can be created out of three variable nodes because 3 edges can be constructed between a pair of variables (i.e., *x* → *y, x* ← *y* or *x* ↮ *y*) and there are three pairs (i.e., {*b, v*}, {*v, s*}, {*s, b*}; 3 × 3 × 3). A total of 6 node combinations can be used for {*y*_*c*_, *y*_*s*_, *y*_*v*_} since we have three (IPL_b1_, pSTG_b3_, pSTG_b5_), two (DLPFC_s3_, Cereb_s5_), and single (aSTG_v5_) candidate brain signals for *b, s*, and *v*, respectively (3 × 2 × 1). Thus, we evaluated which of the 162 (27 edges structures x 6 nodes combinations) possible *G*s is most likely. Since these *G*s differ in complexity, we used the Bayesian Information Criterion (BIC) values for the comparison (the smaller a BIC value is, the more likely a graph is).

### Statistics

When we searched for the brain signals of the BMBU’s latent variables using the searchlight technique, we applied the mixed effect model to enhance statistical power by complementing the noise in the BOLD signals. On the other hand, we did not apply the mixed effect model to analyze the other data. The significance tests were two-tailed except for the searchlight analysis as specified in Table 1. Also, for the time-resolved searchlight analysis, we implemented the multiple-comparison test (the fdr correction)^85^ for each of the fMRI time frames. In the figures summarizing statistical results, all confidence intervals are the 95% confidence intervals of the mean.

## Data and code availability

Data and code will be available after publication.

## Acknowledgement

This research was supported by the Brain Research Program through the National Research Foundation of Korea (NRF) funded by the Ministry of Science and ICT (No. NRF-2021R1F1A1052020), and SNU R&DB Foundation (339-20220013).

---

**Supplementary Fig. 1.**
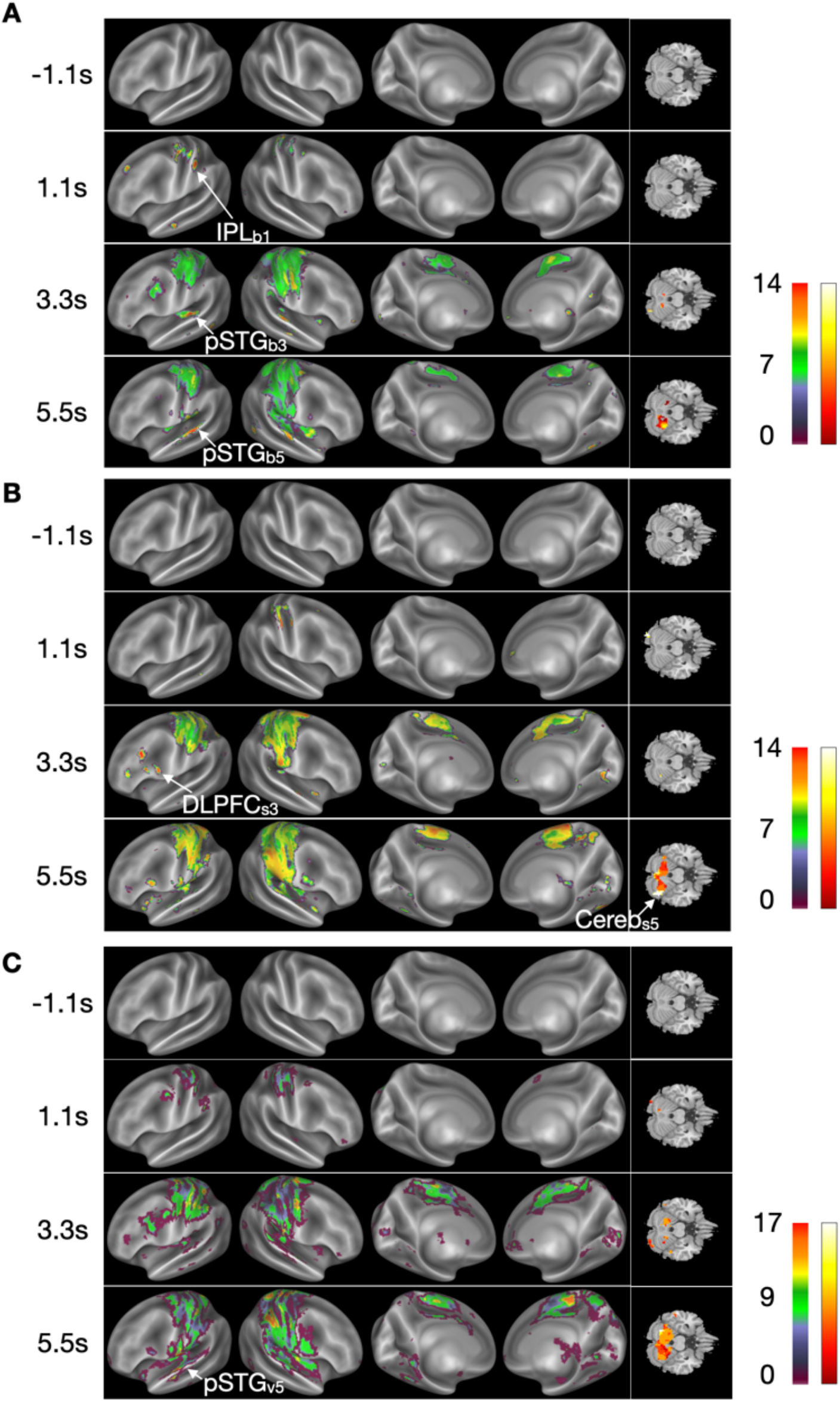
Brain maps of the numbers of the satisfied regressions. **A-C** The number of the satisfied regressions of the candidate brain signals of the inferred class boundary (*b*_(*t*)_; **A**), the inferred stimulus (*s*_(*t*)_; **B**), and the decision variable (*v*_(*t*)_; **C**) on the corresponding regressors (, which are listed in Table 1) are overlaid on the inflated brain hemispheres and the cerebellum of the template brain. The number maps are shown separately for the different within-trial time points relative to stimulus onset. The left and right color bars show the hues corresponding to the number of the satisfied regressions of the maps overlaid on the cortex (left) and the cerebellum (right).

